# Female Dispersion Is Necessary, But Not Sufficient, For The Evolution of Monogamy in Mammals

**DOI:** 10.1101/2020.12.30.424795

**Authors:** R.I.M. Dunbar

## Abstract

Explanations for the evolution of monogamy in mammals typically emphasise one of two possibilities: monogamy evolves when females are overdispersed (such that males cannot defend more than one female at a time) or when males provide a service to the female. However, the first claim has never been directly tested. I test it directly at three levels using data from primates and ungulates. First, I show that the females of monogamous genera do not have territories that are significantly larger, either absolutely or relatively, than those of polygamous genera. Second, using both the Mitani-Rodman and Lowen-Dunbar inequalities, I show that, given their typical day journey lengths, males of most monogamous species could easily defend an area large enough to allow them to monopolise as many as 5-10 females if these ranged solitarily. Finally, I use a model of male mate searching strategies to show that, unlike the males of socially-living polygamous species, the opportunity cost that monogamous males incur is typically more than five times the reproductive success they have by being obligately monogamous. This suggests that the selection pressure dissuading them from pursuing a roving male strategy must be very considerable.

## Introduction

Under classical socioecology theory, females distribute themselves in the landscape in ways that allow them to maximise their fitness by trading off the costs and benefits of group-living (Krause & Ruxton 2000). Males then map themselves onto the distribution of females in an ideal free distribution as they compete for access to the females. Most analyses agree that the ancestral state for mammals, including primates, was some form of dispersed, solitary lifestyle (Müller & Thalmann 2000; Shultz et al. 2011). Although different analyses have argued for alternative routes whereby group-living emerged from this ancestral state (compare Shultz et al. [2011] with Kappeler & Pozzi [2019]), the substantive issue is how males respond to the grouping patterns of females rather than categorical classifications of social systems. The critical contrast lies in whether females live socially in groups or forage alone (with or without dependent young) (Emlen & Oring 1977; see also Rutberg 1983) and whether, when females forage alone, males are forced to choose between attaching themselves to one female in a monogamous relationship or opting for a more promiscuous roving-male strategy

Historically, three principal hypotheses have been proposed as explanations for monogamy in mammals, namely (1) biparental care, (2) female overdispersion and (3) protection against external threats (such as predation or infanticide). This issue continues to attract debate with several recent analyses (Dobson et al. 2010; Shultz et al. 2011; Opie et al. 2013; Lukas & Clutton-Brock 2013; Kappeler & Pozzi 2018) disagreeing with each other. Although a likely explanation in the case of birds (Shultz & Dunbar 2010), biparental care can be excluded for mammals since, the canids notwithstanding, every substantive analysis has agreed that biparental care evolves *after*, or independently of, the adoption of pairbonding (Dunbar, 1995a; Komers & Brotherton 1997; Brotherton & Komers 2003; Opie et al. 2013; Lukas & Clutton-Brock 2013). Beyond that, however, there seems to be little agreement as to whether the main factor selecting for monogamy has been female dispersion, male mating strategies, or other services offered by the male (van Schaik & Dunbar 1990; Komers & Brotherton 1997; Brotherton & Komers 2003; Opie et al. 2013; Lukas & Clutton-Brock 2013; Kappeler & Pozzi 2018).

There are two separate questions here. One is whether, *and* why, females forage alone rather than in groups, and the other is why, given that they do, males are willing to attach themselves to a single female rather than opting for roving male polygamy if this would give them access to many more females. For obvious reasons, females foraging alone in their own territories is a *necessar*y condition for the evolution of monogamy. However, the claim that female overdispersion due to foraging competition is the driver for pairbonding by males seems to be based largely on assumption, since no *nutritional* evidence has ever been adduced to show that monogamous females *need* to forage alone to ensure access to sufficient food. Overdorff & Tecot (2006), for example, reported that, in the lemur *Eulemur rubriventer*, agonistic encounters against both neighbouring conspecifics (rare) and other sympatric lemur species (far more common) occurred more often during the season of food abundance, and concluded from this that resource defence was the main factor selecting for monogamy. Aside from the fact that there is much less to gain from defending resources when they are abundant (Dunbar 1988), this evidence does not conclusively tell us whether fights are over food sources or a simply consequence of adjacent groups coming into contact more often because they converge on good food patches (which, on the evidence given, seems more likely).

In fact, comparative analyses cast doubt on the role of foraging: they suggest that monogamous female mammals do not necessarily live in larger territories than species where females live in groups, even though their densities might be lower (Dunbar 1988; Lukas & Clutton-Brock 2013). More importantly, time budget models for monogamous gibbons (Dunbar et al. 2019) and other medium-sized arboreal primates (Korstjens & Dunbar 2007; Korstjens et al. 2018) indicate that, except on the margins of their biogeographical distributions, there is nothing in their *ecology* to prevent them living in significantly larger social groups than they actually do.

Irrespective of the reasons why females live alone, the crucial question is whether this is also a *sufficient* explanation for pairbonded monogamy. It has been widely suggested that males are forced into monogamy because they cannot defend several females simultaneously against rival males. Brotherton & Manser (1997), for example, concluded that inability to defend a feeding territory could not explain monogamy in dikdik antelope, and suggested that this genus’ monogamous mating system was more plausibly explained as mate-guarding. An alternative version of this hypothesis suggests that males are risk averse and seek to minimise the variance in sirings by opting for the certainty of one female rather than the risk of ending up with no matings at all if they pursue a roving male polygyny strategy (Komers & Brotherton 1997; Brotherton & Komers 2003). Other studies have suggested that males provide a service to the female by minimising the risk of infanticide (e.g. van Schaik & Dunbar 1990; Opie et al. 2013), defending a food supply by keeping conspecifics out of their territory (Williams et al. 2004), or providing protection from predators (Dunbar & Dunbar 1980).

Ever since Hamilton’s (1964) seminal papers, evolutionary biologists have understood that the crucial evolutionary question is not what benefits an animal gets from a particular trait but the balance between these benefits and the opportunity cost, or ‘regret’, that it incurs by not adopting the alternative(s). The magnitude of the difference in benefits between the trait that animals have adopted and the alternative that they have not provides a measure of the magnitude of the selection pressure favouring the former. Too many comparative analyses overlook this and, as a result, risk failing to distinguish between genuine selection factors that have driven evolution and windows of evolutionary opportunity that arise once a trait has been acquired.

In this paper, I focus on testing the hypothesis that monogamy arises because males cannot monopolise mating access to more than one female when females are overdispersed. I do this at three separate levels. First, I ask whether the females of monogamous species really do occupy larger territories than the females of polygamously mating territorial species, either in absolute terms or relative to their metabolic demands (*Hypothesis H1*). Despite its obvious relevance, the latter possibility seems never to have been directly tested. Second, I test whether the males of monogamous species could, in principle, defend territories larger than those they actually occupy, and hence could access more than one reproductive female (*Hypothesis H2*). Finally, I test whether, in terms of fertilisations obtained, it would pay males to abandon spatial localisation altogether and adopt a roving male strategy that allows them to search for females without necessarily having to defend a territory (*Hypothesis H3*).

I test these hypotheses on the 12 genera of monogamous anthropoid primates (mainly hylobatids and the New World callitrichids and cebids), one nominally monogamous prosimian (*Lepilemur*, whose social system has been described as “dispersed monogamy”: Hilgartner et al. 2012) and two monogamous miniature antelope (one a facultative, the other an obligate monogamist: Dunbar & Dunbar 1980; Adamczak & Dunbar 2008). The callitrichids are somewhat anomalous because, although they have traditionally been considered to be monogamous, their mating systems are in fact more complex and variable, with individual groups switching between monogamy, polygamy, polyandry and polygynandry over time (Goldizen 1988; Dunbar 1995a,b; Digby et al. 2007). Nonetheless, I include them in the monogamous sample because they normally have only a single breeding female, irrespective of the composition of the social group, due to puberty being suppressed in maturing females (Abbott 1984). For comparison, I use data from two genera of territorial polygamous monkeys (*Colobus* and *Cercopithecus*) and one non-territorial polygamous ape (*Gorilla*) whose social systems can best be defined as area- or group-defence polygamy (these sometimes being indistinguishable). Finally, as benchmarks for the models, I include data on 11 genera of nocturnal prosimians that have a roving male mating system and one genus of multimale-grouping anthropoids (*Papio*) that live in very large, stable social groups. If the models are empirically valid, they should consistently predict that the prosimian males will prefer a roving male strategy and that the *Papio* males should always prefer to be social.

Rather than focus on species averages, as most comparative analyses do, I focus on individual populations on the grounds that animals’ behaviour is a response to local ecological and demographic conditions rather than rigidly inherited species-typical traits (see Dunbar 1995b; Dunbar et al. 2009). Although comparative studies typically assume that species have a characteristic social system, in fact all primate social systems can be remarkably flexible. Almost all monogamous primates and ungulates, for example, occasionally have groups with more than one breeding female (Kappeler & Pozzi 2018; Adamczak & Dunbar 2008). Similarly, although the typical social group of all three of the comparator polygamous genera is a unimale group, in fact multimale groups occur in all three genera (Fashing 2007; Enstam & Isbell 2007; Robbins 2007). *Cercopithecus* provides a particularly relevant test case because, although its basic social and mating system is a one-male harem group, some species (notably *Cercopithecus mitis*) are subject to temporary influxes of males when there are many females simultaneously in oestrus in the group (Henzi & Lawes 1987; Cords 2004; Roberts et al. 2014; Gao & Cords 2020). This variability is important for present concerns because it implies that animals can, and do, vary their social and mating strategies in response to local environmental and demographic contexts. My concern is to determine the circumstances under which such switches of socio-sexual strategies might occur in populations of the different genera, and to use that as a means of gaining insight into the selection pressures acting on individual taxa.

## Methods

### Data

I sourced data on day journey length, territory size, mean social group size, mean number of females per group, group density (groups per km^2^) and interbirth interval (in days) from Campbell et al. (2007), with updates from a search of the post-2007 primary literature. I supplemented these with data on *Gorilla* from Doran & McNeilage (1998) and Lehmann et al. (2008), and Doran-Sheehy & Boesch (2004). Data for gibbons are taken from Dunbar et al. (2019). Altogether, 90 populations from 48 species representing 15 genera are represented in the haplorrhine sample, with a mean of 5.6±2.8 (range 2-11) populations per genus. In most cases, data are averages from several groups living in the same location. Data on interbirth intervals are used only if they derive from wild populations. Where possible, the value given for each specific study site is used. Data for the nocturnal prosimians were sourced as species means from Campbell et al. (2007) and other primary sources. Data for nine *Papi*o populations were sourced from Dunbar (1992), Hill et al. (2000), Bettridge et al. (2010) and Dunbar et al. (2018a). Field studies of antelope almost never provide data on day journey length. However, I was able to source data for four populations of monogamous miniature antelope (three for klipspringer, *Oreotragus oreotragus*, and one for the socially more variable oribi, *Ourebia ourebi*).

Aside from the gorilla, the species included in these analyses do not vary significantly in body mass. Excluding gorilla, mean body mass is 2.84±3.19 kg for the monogamous species and 5.56±2.59 kg for the comparator polygamous anthropoids (t_12_=-1.33, p=0.346). There is, thus, no reason to correct for body mass in respect of any ecological variables other than those where body mass forms part of the variable of interest.

The data are provided in the *SI Dataset*.

### Analyses

I tested the first hypothesis by comparing mean absolute territory size. I also calculated relative territory size as size per female metabolic body weight (km^2^/kg0.75).

To test the second hypothesis, I determined the maximum territory size that a male could defend, and then estimated the number of females that could live in that territory given the observed mean size of female territories in that particular habitat. To determine the maximum size of territory that a male could defend, we can exploit two alternative indices of territory defendability in primates based on slightly different assumptions about how males detect intruders. Although conceptually simple, both indices predict whether primate species are territorial or not with surprising accuracy.

Mitani & Rodman (1979) showed empirically that primate species are territorial so long as:

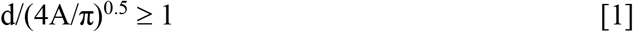

where *d* is the day journey length (in km), *A* the home range area (in km^2^), and the territory is assumed to be a perfect circle. In effect, males can defend a territory against intruders so long as the distance they travel each day while foraging exceeds the diameter of their home range (i.e., they can traverse their territory at least once a day). The ratio of 1 has no particular significance: it is simply an empirical observation that differentiates most territorial from most non-territorial species. It probably reflects something about the frequency with which a male can check out his territory if he travels randomly around it, given his observed daily travel: in effect, it tells us something about the mean interval between successive visits to a particular location during which an intruder would have unfettered access to the resources at that location. We can determine the maximum size of territory the male could defend, *A_max_*, by inverting Eq. [1] and setting it equal to 1 to obtain:

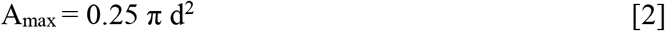

This simply assumes that if some species can defend larger ranges, there is in principle no reason why all species cannot do so.

Lowen & Dunbar (1996) reformulated the Mitani-Rodman inequality on the assumption that the critical issue is how often a male can check the boundary of his territory rather than the whole area enclosed within this boundary (as the Mitani-Rodman inequality assumes). Using the Maxwell-Boltzman gas dynamics equation, they found, again empirically, that primate species are territorial whenever:

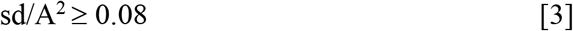

when the arc of the territory boundary, *s*, within which a male can detect an invading rival is taken to be s=0.5 km (i.e. 250 m either side of the point at which the male arrives at the boundary of his territory). Lowen & Dunbar (1996) noted that, while providing an improvement in the accuracy with which territorial species were differentiated from non-territorial species, this formulation mainly serves to confirm the validity of the Mitani-Rodman inequality. However, the value of *s* (the width of the boundary detection arc) is crucial, since it affects how much of the territory’s circumference the male can monitor on any given visit, and hence how long it will take him to check the whole perimeter while travelling randomly about his territory (in effect, the time between successive visits to the same location). As *s* decreases, the criterion on the right hand side increases proportionately, and correspondingly decreases as *s* increases. Subject to this constraint, we can determine the maximum defendable territory size in exactly the same way as we did with the Mitani-Rodman inequality, by inverting Eq. [3] and setting it equal to 0.08*(0.5*s*^−1^):

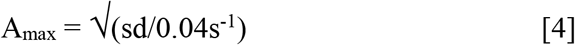

Following van Schaik & Dunbar (1990) and Dunbar (1995a), we can use Eqs. [2] and [4] to calculate the number of reproductive females a male could expect to include within his maximally defendable territory by dividing *A_max_* (less *A_male_*, the share of the observed territory that the male needs for his nutritional requirements) by the size of the observed territory required to support one female and her offspring (*A_female+young_*). The number of females a male can expect to monopolise, *E*(*f*), is then given by:

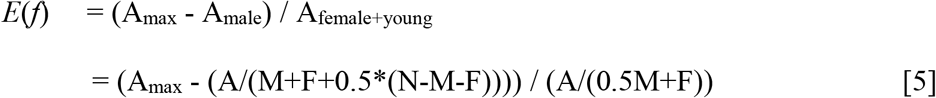

where *M* is the number of males in the group (in our sample, *M*=1 in all cases except *Papio*), *F* is the mean number of females per group and *N* is the mean size of the group. For computational convenience, dependent young are counted as the energetic equivalent of half an adult on average. Except for *Gorilla*, males and females are treated as being of equivalent body mass. Because *Gorilla* are the most sexually dimorphic of all primates, I weight males as equivalent to two females in their case.

The two indices are highly correlated in their prediction of the maximum size of territory that a male could defend (for the 79 populations in the sample for which there are day journey length data, *r*=0.949, p≪0.0001). Nonetheless, compared to the Mitani-Rodman index, there is a consistent tendency for the Lowen-Dunbar index to overestimate the defendable territory size when these are small and underestimate them when they are large. The exact numerical values differ depending on the value of *s* in Eq. [4]. The consequence of this is best illustrated by the effect it has on the number of females that the male could defend access to. Compared to the Mitani-Rodman index, the Lowen-Dunbar index underestimates the maximum number of defendable females when *s* is small (Fig. S1a) and overestimates it when *s* is large (Fig. S1c). The two indices converge at *s*≈0.2 (Fig. S1b). Although Lowen & Dunbar (1996) used a detection arc of *s*=0.5 km, a value of *s*=0.2 (i.e. 100 m either side of the male) is probably more realistic limit for animals that live in forest, as all the genera in our sample except *Papio* do. Because of the uncertainty surrounding *s* and the fact that the Mitani-Rodman index is always more conservative for species that have small day journeys and territories, I use this index in preference, but the equivalent results for the Lowen-Dunbar inequality are given in the online SI.

For perhaps obvious reasons, data on day journey length are not available for any of the nocturnal prosimian species. I therefore used the observed mean territory sizes for each of the sexes and, given that they are pursuing a roving male strategy, assumed that the males are defending the largest territories they can. At worst, we can be sure that they cannot do worse than they already actually do. For each species, I calculated *E*(*f*) as male territory size divided by female territory size (less the male’s share estimated as a third).

I then calculate the selection ratio of the two strategies, so that the male’s decision rule should be:

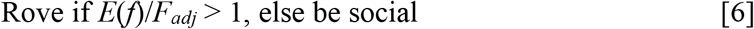

where *F_adj_* is the mean number of females living together in a social group adjusted for the genus’s reproductive rate (one in the case of monogamous genera, and the mean observed number of females in the group for polygamous genera). Because most callitrichids twin and some produce two litters a year, I use the genus-specific number of infants born to a monogamous male as the social baseline in these cases. This is the equivalent of four females for *Callithrix*, *Saguinus* and *Cebuella* (whose females habitually produce twins twice a year) and two for *Callimico* (who produce one infant twice a year: Porter et al. 2001) and *Leontopithecus* (who produce twins once a year: Bales et al. 2001). Since all these callitrichid genera have biparental care, I assume, following Dunbar (1995a), that, when males rove, the females will be obliged to revert to the ancestral anthropoid state of producing a singleton offspring once a year. The magnitude of ratio [6] is a measure of the strength of selection favouring roving.

The third hypothesis asks whether, in terms of actual numbers of offspring sired, a male would be better off abandoning his territory and instead simply roaming as widely as he can in search of females. To calculate the number of sirings such a male could expect to obtain, I used a model originally developed for feral goats (Dunbar et al. 1990) and later extended to great apes (Dunbar 2000, 2001). The model assumes that a male searches randomly without geographical limit, but subject to his own daily ranging limits. The number of progeny, *p*, that a roving male can expect to sire during the average female reproductive cycle, *E*(*p*), is:

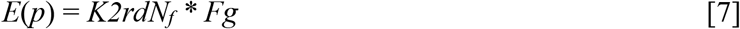

where *K* is the average interbirth interval for the population (reproductive cycle length, in days), *r* is the detection distance, *d* the day journey length (in km) (so 2*rd* is the search path that the male carves out each day as he forages), *N_f_* the density of female groups (groups/km^2^, defined as *A*-1 assuming, conservatively, no overlap in territories), *F* the average number of adult females in a female group and *g* is the probability that a female will be in oestrus on any given day. This model predicts with almost no error the proportion of males that are social (i.e. are found associating with groups of females on any given day) in populations of chimpanzees, gorillas, orang utans and humans (Dunbar 2000), as well as feral goats (Dunbar et al. 1990). Most importantly, the regression line for both the model predictions and the data pass through the point of equilibration where males should be undecided whether or not to be social because the payoffs are equal (see Fig. 1 in Dunbar 2000). This indicates that, notwithstanding its simplifications and notwithstanding the fact that it ignores both the number and behaviour of rival males, the model correctly predicts what males should do across a range of mammalian genera.

Eq. [7] is made up of two components: the frequency with which a roving male can expect to encounter a group of females each day as he travels about his territory, *K2rdN_f_*, and the expected number of fertilisable females in the group when he does find one, *Fg.* The first component is adapted from the Maxwell-Boltzmann gas dynamics equation and consists of two parts: the probability that a randomly searching male will encounter a female group during an average reproductive cycle (calculated as the area that he can search during the course of a reproductive cycle of length *K* days, given a daily search path of *d* km length and a detection distance either side of his path of travel of *r* km width) and the density of female groups, *N_f_*. In the light of Fig. S1, I use 2*r*=0.200 km. It does not matter whether a male searches the same locations repeatedly as the female groups are also mobile, and so have a random chance of occupying a space that the male has previously searched by the time of his next visit. The second part of Eq. [6] gives the mean number of females in oestrus (i.e. that are fertilisable) that a male can expect to find in a group of *F* females. Primate females are typically receptive for about 5 days on each of three successive menstrual cycles during any given reproductive cycle, hence *g*=15/*K*. The model assumes that female reproductive cycles are not in synchrony, so that females become available for mating at random. As before, the ratio of the payoff for a roving male to that for a social male, *E*(*p*)/*F_adj_*, is an index of the selection pressure favouring roving, as in inequality [6].

**Fig. 1.**
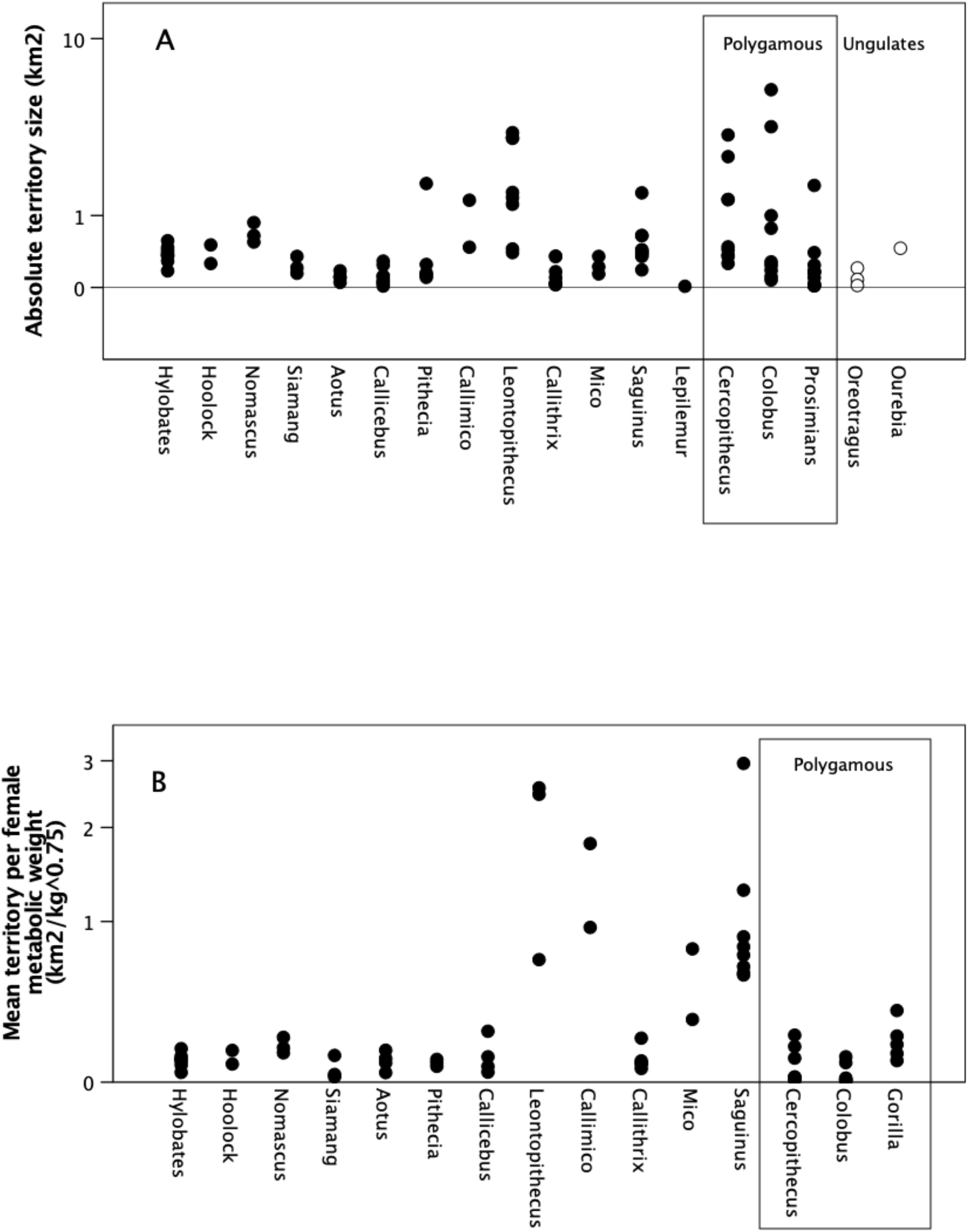
(a) Observed absolute territory size (km^2^) and (b) relative territory size per female adjusted for metabolic weight (km^2^/kg^0.75^). *Gorilla* not included in (a) because of its markedly larger body size. Unfilled symbols: ungulates.

### Phylogenetic analysis

It is conventional to apply phylogenetic controls when undertaking comparative analyses, so as to avoid the problem of inflated sample sizes when closely related species share traits by descent from a common ancestor. Phylogenetic methods are not used in the present analyses for four reasons. First, none of the analyses test functional (i.e. causal) hypotheses that compare correlated changes on two dimensions: I simply ask how a given taxon performs on a particular behavioural dimension. There are *no* phylogenetic methods available for one-dimensional comparisons of this kind, even if this was desirable (see Dunbar et al. 2018b). Second, most of the analyses are explicitly *within-population* comparisons: they compare alternative outcomes for a particular species at a particular location. Phylogenetic inertia cannot be an issue in such cases, especially as the variance in most of the variables of interest are subject to strong environmental and demographic influence (Dunbar et al. 2009). Third, all the comparisons presented here are at the within-genus level. Since most of the variance in primate behaviour and anatomy lies at genus rather than species level (Harvey & Clutton-Brock 1985), genus-level analyses effectively control for phylogenetic effects. I therefore follow Dunbar et al. (2009) and treat species within a genus as ecological populations whose behaviour is driven by environmental conditions rather than biological constraints. Finally, notwithstanding to two previous points, the phylogenetic signals for all behavioural and ecological measures for primates are close to *λ*=0 (Kamilar & Cooper 2013). Indeed, for no comparative analysis of primate behavioural data so far reported has the use of phylogenetic methods yielded results that differ significantly from analyses that do not use phylogenetic methods (or, more importantly, produce non-significant results where uncorrected analyses are significant).

## Results

### Comparative territory size (H1)

I tested whether monogamous genera have larger territories than polygamous genera in two ways. Fig. 1a plots absolute territory size for the monogamous genera and the comparator polygamous genera, as well as the two monogamous browsing antelope (on the right hand side). *Gorilla* is not shown because its body mass is approximately two orders of magnitude larger than any of the other taxa. It is obvious that, with the exception of *Leontopithecus*, the monogamous genera do not have territories that are especially large by comparison with those of primates that live in polygamous territorial groups. Indeed, many of the monogamous genera, notably the cebids, have much smaller territories than the polygamous genera. Even monogamous antelope do not seem to differ in this respect (Fig. 1a). Brotherton & Manser (1997) also found that there was no difference between monogamous and polygamous groups of dikdik antelope (genus *Madoqua*) in either the size or the resource quality of territories.

It may be that the females of monogamous genera are obliged to live in territories that are *relatively* larger, given the energy demands of reproduction and the number of young they have to support. Fig. 1b tests this by plotting territory size standardised to female metabolic mass (km^2^/female*kg^0.75^). *Gorilla* is included in this graph. While *Callimico*, *Leontopithecus* and *Saguinus* have *relative* territory sizes that are larger than other genera, in general the monogamous species do not differ, as a set, from the polygamous genera. In other words, with the exception of three of the four callitrichid genera, none of the monogamous genera are forced to be overdispersed by the high energy demands of their females or the poor quality of their territories.

On balance, then, there is no compelling evidence to support the suggestion that monogamous species have larger absolute or *per capita* territories than territorial polygamous species (Hypothesis 1).

### Maximally Defendable Territory Size (H2)

To determine whether males in monogamous genera are obliged to associate with one female because they cannot defend a larger territory, I first used the Mitani-Rodman inequality (Eq. [2]) to calculate the maximum size of territory that a male could defend, and then determined (using Eq. [5]) how many females could be encompassed within this territory. To account for the fact that genera differ in their baseline number of females (i.e. the actual number of females in a group), I rescaled each population’s value as a ratio of the observed mean number of females in its groups, so that all populations have the same baseline (with the callitrichids adjusted by relative fertility of the females).

Fig. 2 plots the predicted number of females that a territorial male could expect to have within its maximally defendable territory as a ratio of the baseline number he would have if he was social (i.e. lived with his group). The dashed line demarcates a ratio of 1 where the payoffs of the two strategies would be equal. Males should prefer a roving strategy if the payoff ratio is >1, but prefer being social if the ratio is <1. The model correctly predicts that *Papio* should prefer to be social (mainly because of the large size and low density of its groups) and that all the nocturnal prosimians should prefer the roving male strategy. The model thus makes the correct predictions for our benchmark taxa, confirming its reliability.

**Fig. 2.**
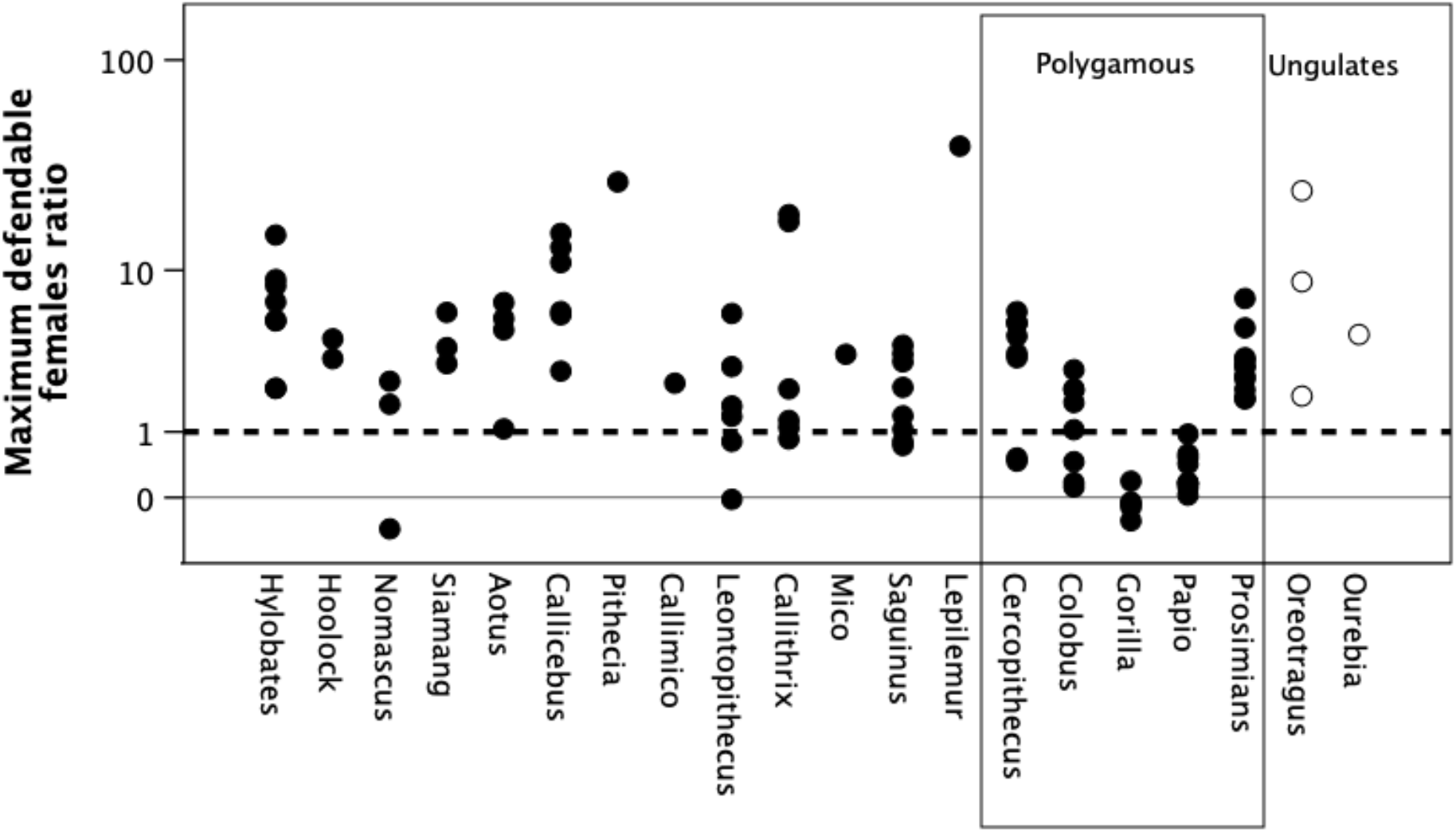
Ratio of maximum number of defendable females, as predicted by the Mitani-Rodman equation, to the default baseline of observed mean number of females per group, for individual populations. The baseline is one female for all monogamous taxa, except *Callithrix, Mico* and *Saguinus* for whom the equivalent baseline is four females (pairbonded females produce twins twice a year) and *Callimico* and *Leontopithecus* for whom it is two females (see text). For the polygamous genera, the mean number of females per group is 7.1 for *Cercopithecus*, 3.8 for *Colobus* and 4.4 for *Gorilla*, 1.0 for the prosimians and 16.5 for *Papio.* When the ratio = 1 (heavy dashed line), the payoffs are equal and males should be ambivalent about their preferred strategy. Unfilled symbols: ungulates.

I use a one-sample t-test to determine whether each genus in Fig 2 differs from a baseline value of one. The tests are one-tailed positive because we are only interested in whether males do better by roving than by being social; by definition, a significant negative result (a male does significantly worse by roving) is as much evidence against the hypothesis as a non-significant result. In most cases, males would be able to monopolise access to significantly more females by pursuing a roving strategy (Table 1). For the monogamous genera as a set, the distribution of *p*-values is more significant than would be expected by chance (Fisher’s meta-analysis: χ^2^=63.21, df=2*9=18, p≪0.0001), indicating a consistent underlying pattern. *Nomascus* and *Leontopithecus* are the least significant (both occupy marginal or degraded habitats, and have absolutely larger territories than the other genera in their respective taxonomic families: Fig. 1a), with *Callithrix* on the margin. Even so, in at least some populations of even these genera, it would pay males to be polygamous. Among the comparator genera, *Cercopithecus* would benefit by roving, but *Colobus* and *Gorilla* would not. That *Gorilla* males would always do significantly better by being social and staying with a group of females reflects the costs of travel for great apes and their short day journeys (Lehmann et al. 2008a,b).

**Table 1.**
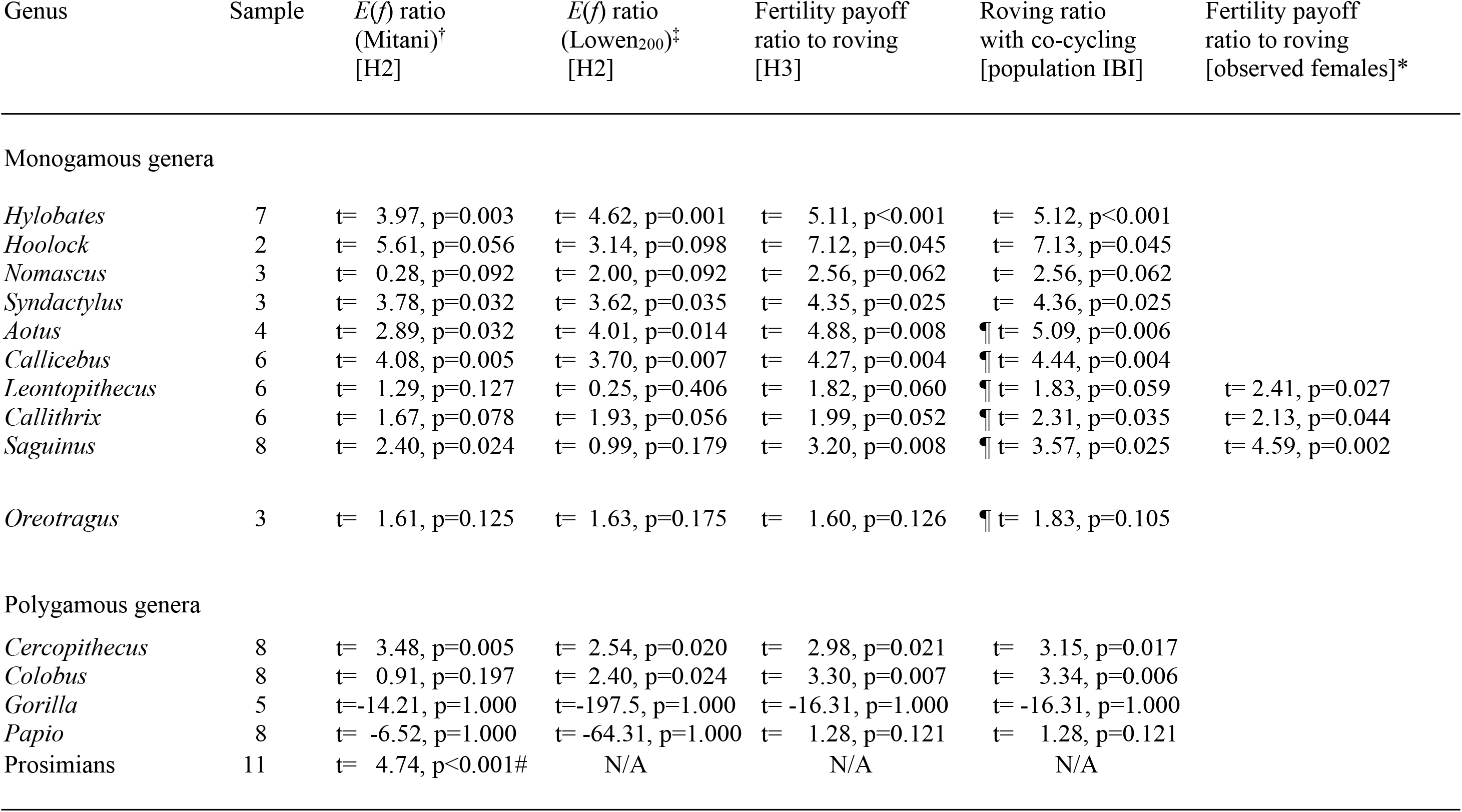
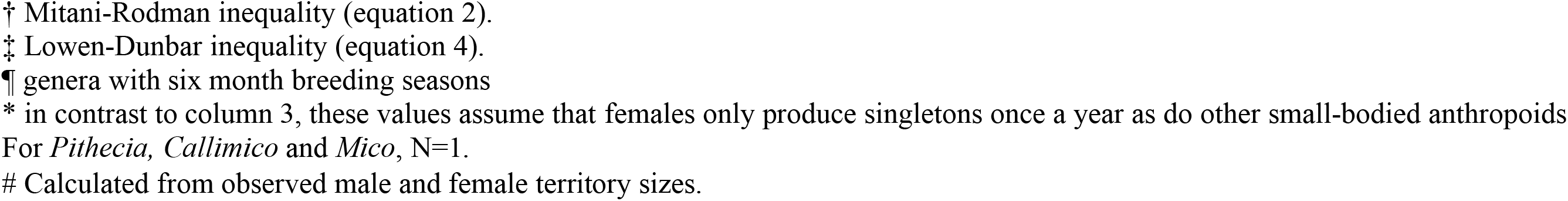
One sample t-tests against H_0_=1 for each of the main indices for each genus for Hypotheses H2 and H3

| Genus | Sample | $E(f)$ ratio<br>(Mitani)<sup>†</sup><br>[H2] | $E(f)$ ratio<br>(Lowen<sub>200</sub>)<sup>‡</sup><br>[H2] | Fertility payoff<br>ratio to roving<br>[H3] | Roving ratio<br>with co-cycling<br>[population IBI] | Fertility payoff<br>ratio to roving<br>[observed females]* |
| --- | --- | --- | --- | --- | --- | --- |
| Monogamous genera |  |  |  |  |  |  |
| *Hylobates* | 7 | t= 3.97, p=0.003 | t= 4.62, p=0.001 | t= 5.11, p<0.001 | t= 5.12, p<0.001 |  |
| *Hoolock* | 2 | t= 5.61, p=0.056 | t= 3.14, p=0.098 | t= 7.12, p=0.045 | t= 7.13, p=0.045 |  |
| *Nomascus* | 3 | t= 0.28, p=0.092 | t= 2.00, p=0.092 | t= 2.56, p=0.062 | t= 2.56, p=0.062 |  |
| *Syndactylus* | 3 | t= 3.78, p=0.032 | t= 3.62, p=0.035 | t= 4.35, p=0.025 | t= 4.36, p=0.025 |  |
| *Aotus* | 4 | t= 2.89, p=0.032 | t= 4.01, p=0.014 | t= 4.88, p=0.008 | ¶ t= 5.09, p=0.006 |  |
| *Callicebus* | 6 | t= 4.08, p=0.005 | t= 3.70, p=0.007 | t= 4.27, p=0.004 | ¶ t= 4.44, p=0.004 |  |
| *Leontopithecus* | 6 | t= 1.29, p=0.127 | t= 0.25, p=0.406 | t= 1.82, p=0.060 | ¶ t= 1.83, p=0.059 | t= 2.41, p=0.027 |
| *Callithrix* | 6 | t= 1.67, p=0.078 | t= 1.93, p=0.056 | t= 1.99, p=0.052 | ¶ t= 2.31, p=0.035 | t= 2.13, p=0.044 |
| *Saguinus* | 8 | t= 2.40, p=0.024 | t= 0.99, p=0.179 | t= 3.20, p=0.008 | ¶ t= 3.57, p=0.025 | t= 4.59, p=0.002 |
| *Oreotragus* | 3 | t= 1.61, p=0.125 | t= 1.63, p=0.175 | t= 1.60, p=0.126 | ¶ t= 1.83, p=0.105 |  |
| Polygamous genera |  |  |  |  |  |  |
| *Cercopithecus* | 8 | t= 3.48, p=0.005 | t= 2.54, p=0.020 | t= 2.98, p=0.021 | t= 3.15, p=0.017 |  |
| *Colobus* | 8 | t= 0.91, p=0.197 | t= 2.40, p=0.024 | t= 3.30, p=0.007 | t= 3.34, p=0.006 |  |
| *Gorilla* | 5 | t=-14.21, p=1.000 | t=-197.5, p=1.000 | t= -16.31, p=1.000 | t= -16.31, p=1.000 |  |
| *Papio* | 8 | t= -6.52, p=1.000 | t= -64.31, p=1.000 | t= 1.28, p=0.121 | t= 1.28, p=0.121 |  |
| Prosimians | 11 | t= 4.74, p<0.001# | N/A | N/A | N/A |  |

† Mitani-Rodman inequality (equation 2).
‡ Lowen-Dunbar inequality (equation 4).
¶ genera with six month breeding seasons
\* in contrast to column 3, these values assume that females only produce singletons once a year as do other small-bodied anthropoids For *Pithecia*, *Callimico* and *Mico*, N=1.

Fig. 2 also plots the number of defendable females given by Eq. [5] for the two genera of monogamous miniature antelope (klipspringer, *Oreotragus*, and the oribi, *Ourebia*). In both cases, males could easily defend territories large enough to monopolise access to 5-10 females (the size of most harem groups in medium-sized antelope). Yet, like most of the miniature antelope, these species are monogamous, and especially so in the case of *Oreotragus*. *Oreotragus* is the one antelope species that comes closest to primate levels of social attentiveness and bondedness: the modal separation between the male and female is just 3-5 m and the pair constantly monitor each other, with one moving or resting when the other moves or rests (unpublished data; see also Dunbar & Dunbar 1980; Dunbar & Shultz 2010).

A possible confound is that Eq. [5] assumes that the habitat is completely packed with female territories. If the density of territories (i.e. groups) is significantly less than that assumed by continuous packing, then polygamy might not be a viable strategy. Fig. 3 plots the mean density of groups for the monogamous taxa, based on actual field estimates. The median observed group density (indicated by the solid line) is 3.30 groups per km^2^; the median estimated density based on Eq. [5] assuming no overlap between the ranges of neighbouring females is 2.91 groups per km^2^ (indicated by the dashed line). Although some genera (such as *Callithrix*) have higher densities and some (such as *Leontopithecu*s) have lower, on average actual density is not consistently lower than the maximum packing assumed by the model. This makes it very unlikely that Fig. 2 seriously overestimates the payoff to promiscuous males.

**Fig. 3.**
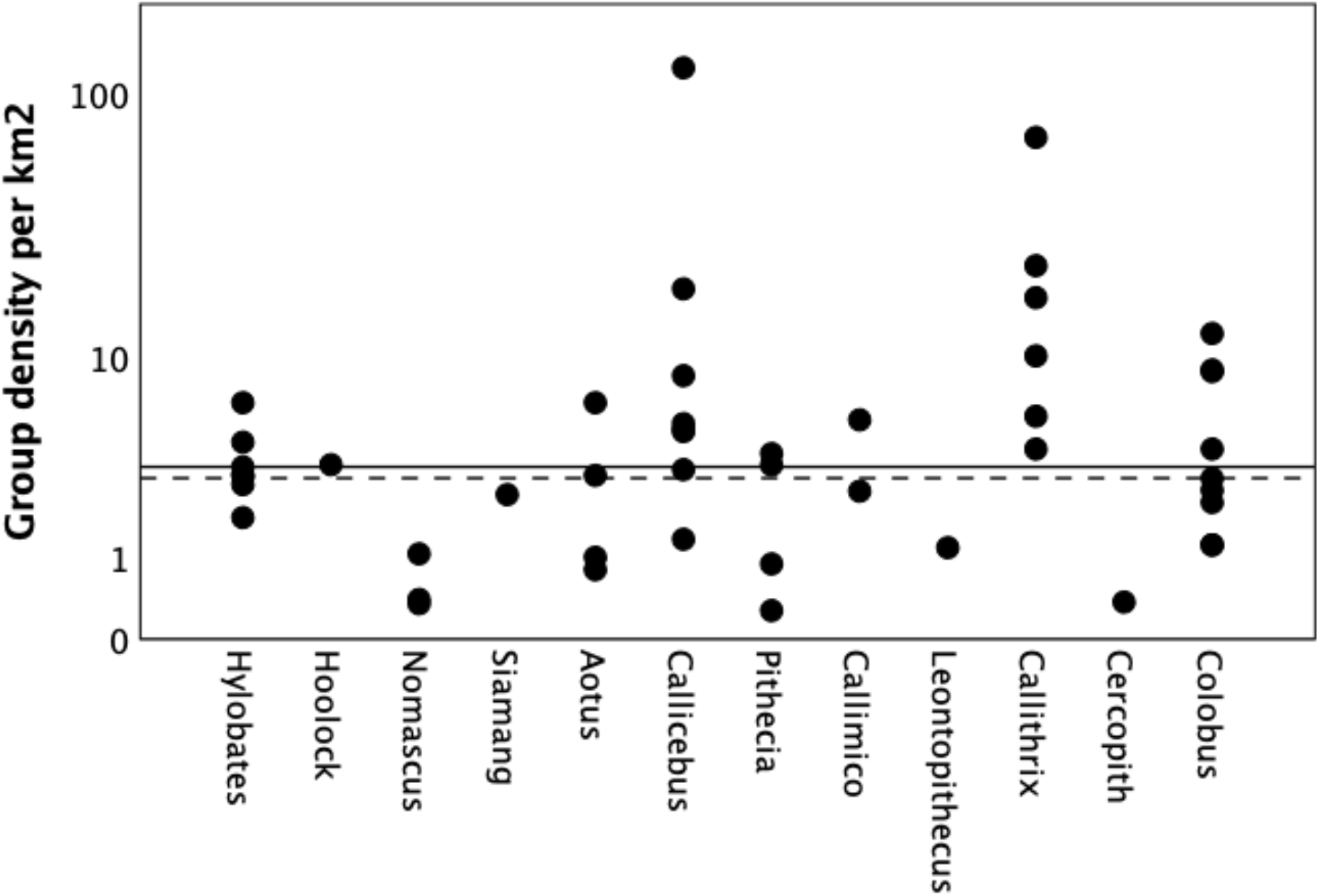
Observed density of groups for monogamous genera, for individual populations. The solid line indicates the median value for the observed data; the dashed line indicates the median value estimated by the model (based only on day journey length and assuming no overlap of adjoining territories). The model does not overestimate the density of groups.

Fig. S2 plots the equivalent results for the anthropoid primates using the Lowen-Dunbar inequality (Eq. [4]). Since night journey length data are not available for any of the prosimians, it is not possible to calculate the maximally defendable territory size for these taxa, but the model correctly predicts that *Papio* males should prefer to be social. The results across individual populations are significantly correlated with those in Fig. 2 (r=0.843, N=96, p<0.0001 1-tailed). The distribution of datapoints across genera is broadly similar to that for the Mitani-Rodman inequality, though *p*-values are lower in some, but not all, cases (Table 1).

Taken together, these results suggest that even though some populations of the monogamous genera would do better by being monogamous, most would be significantly better off pursuing a roving male strategy and this is as true for antelope as it is for primates. In sum, Hypothesis 2 does not receive convincing support, at least as a universal explanation for monogamy even if it might explain some cases.

### Fertility payoff to roving (H3)

Merely being able to control a large territory containing many females does not guarantee higher fitness. What is important is the chances that the females will be in oestrus when the male finds them. With interbirth intervals as long as they are in primates (up to 39 months in the case of gibbons), a randomly searching male is very likely to chance on most females when they are not in oestrus. I determined the fitness payoff to a roving male using Eq. [7] to estimate how many females he would locate when they were actually in oestrus. The time base is the interbirth interval for the population, so that social males would sire exactly one offspring for each female in their group. Fig. 4 plots the payoff ratio to a roving male, *E*(*p*)/*F_adj_*, in terms of the expected number of sirings he would gain relative to that obtained by a social male for each sampled population (adjusted, as in Fig. 2, for the higher reproductive rates of the callitrichids). The dashed line corresponds to a payoff ratio of. As with Fig. 2, males should prefer a roving strategy if the payoff ratio *E*(*p*)/*F_adj_* >1, but prefer to be social if *E*(*p*)/*F_adj_* ≤1. There are no day journey data for nocturnal prosimians, so no predictions are possible for these populations, but the model correctly predicts that *Papio* would prefer to be social.

**Fig. 4.**
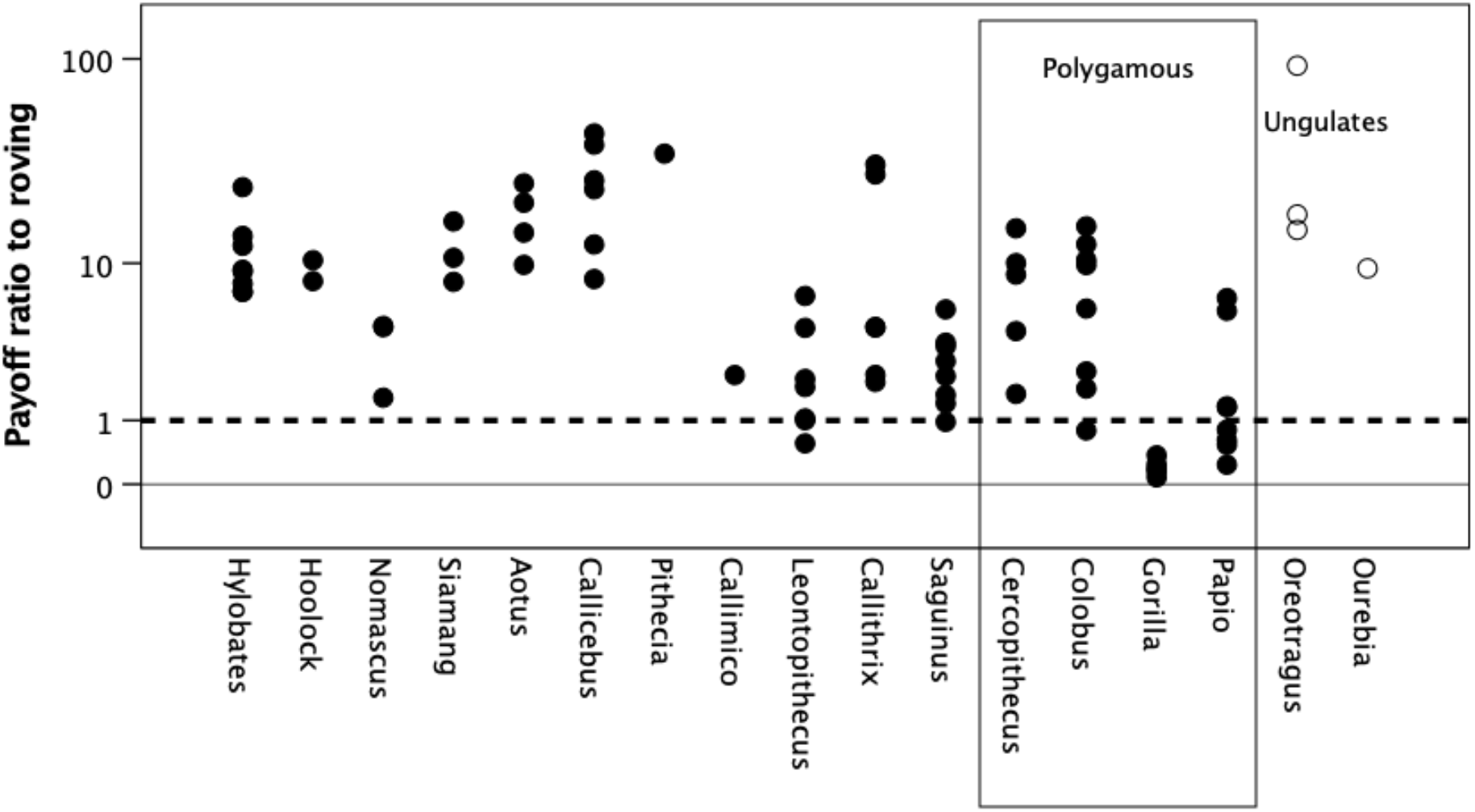
Payoff ratio to promiscuous males (‘roving male strategy’) predicted by the male mating strategies model, for individual populations. The payoff ratio is the number of offspring sired during the average population-specific reproductive cycle (i.e. interbirth interval) by a promiscuous roving male divided by that expected for a social male (which equals female group size, as in Fig. 2). The dashed line at a ratio of 1 indicates the point of equilibrium where the payoffs are equal and males should be ambivalent about which strategy to adopt. When the ratio lies above this line, males would do better by opting for roving male polygamy; when it falls below, they would do better by opting to be social (i.e. permanently attaching themselves to a group of females). Unfilled symbols: ungulates.

Although the males in some individual monogamous genera should prefer to be social, most of the genera would benefit by opting for a roving strategy (payoff ratios significantly >1) (Table 1). Among the monogamous genera, only *Nomascus*, *Callithrix* and *Leontopithecus* are not significant (and then only marginally). Even so, the majority of their populations would benefit by roving. Both ungulate genera would also clearly benefit by roving, although the high variance in these data makes the difference not significant. As before, *Gorilla* would clearly benefit by being social, but both *Cercopithecus* and *Colobus* males would be significantly better off roving.

I reran this analysis for detection distances of 2*r*=0.05 km and 2*r*=0.5 km. Although the payoff ratios are slightly smaller or larger, respectively, the conclusions are broadly the same (Fig. S3). With *r*=0.5, even *Colobus* are significantly above *E*(*p*)/*F_adj_*=1; at *r*=0.05, many more genera are not significant (Fig. S3a). It would, however, be necessary to reduce detection distances below *r*=0.02 (10 m either side of the male) to remove the advantage of roving for all the monogamous genera.

Brotherton & Manser (1998) suggested that the problem might be that when females are overdispersed, males are unable to prevent rivals mating with at least some females if two or more are in oestrus simultaneously; as a result, males might opt for monogamy as a risk-averse strategy (the bird-in-the-hand strategy). We can determine how risky roving is using the binomial expansion:

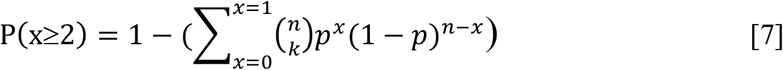

where *P*(*x*≥2) the probability that there will be two or more females simultaneously in oestrus within the area the male can sweep out during a reproductive cycle, *x* is the number of females who are co-cycling, *n* is the total number of females in the area he can search during a full reproductive cycle, and *p* is the probability that any one of them will be in oestrus on any given day (i.e. *p*=15/*K*). Mean *P*(*x*≥2) is 0.056±0.018 across the 71 sampled populations (Fig. S4). Adjusting the payoff to rovers by Eq. (7) yields results that correlate significantly with those from Fig. 4a (r=0.982, N=71 populations, p<0.0001), with a marginal effect on payoff ratios (Table 1: compare Fig. 4 with Fig. S5). Note that *Hoolock* fails to reach significance mainly because its very small sample size (N=2) results in a very large variance estimate.

Some genera in this sample (mainly platyrrhines) are seasonal breeders, although even their mating seasons typically span as much as six months of the year. Since this is likely to increase the chances of females in different groups being in oestrus simultaneously, I reran Eq. [7] with *p*=15/183 days (6 months). Adjusting the payoff to rovers by the resulting probability of co-cycling has a relatively small effect on the significance of the payoff ratios for all genera (Fig. S7). Note, however, it is only the starred genera in Table 1 for whom this is an issue: the other genera do not have breeding seasons.

In sum, with the consistent exception of *Nomascus*, *Leontopithecus* and, marginally, *Callithrix*, the males of all the monogamous genera would do better to adopt a roving male strategy. Once again, even though some populations in these three genera might prefer to be social, this is by no means true for all their populations. Taken as a whole, then, Hypothesis H3 is not supported.

## Discussion

All male mammals face a choice between being social (i.e. living with a group of one or more females) or pursuing a roving male strategy that might give them access to many more females. All three sets of analyses reported here suggest that the males of a wide range of monogamous primate genera would in fact do better by roving. More importantly, at least two genera of monogamous ungulates display similar effects, and do not differ from the primates in the indices I examined. Contrary to conventional wisdom, comparisons with territorial genera where males adopt either area-defence or group-defence polygyny suggest that there is no ecological reason why the males of monogamous genera could not defend larger territories that give them access to more females. This implies either that these males are foregoing considerable additional mating benefits because monogamy offers them some other benefit or that phylogenetic inertia has forced them to retain an ancestral state of monogamy despite the fact that it is maladaptive. Since phylogenetic inertia is not a realistic explanation given that four very distantly related lineages are monogamous, the implication is that monogamy has alternative fitness advantages for males. These benefits must also be advantageous for the female, otherwise she would be not willing to tolerate the male’s continued presence – and, perhaps more importantly, would not be willing to undergo the evolution of the expensive cognitive and behavioural traits associated with pairbonding (Dunbar & Shultz 2021).

It is important to be clear about the logic of these analyses. The approach I adopt is a reverse engineering one that examines the decision that males face in the light of the costs and benefits of pursuing alternative strategies, where these costs and benefits are determined by the lifehistory and social demography of the females *at the actual location where they live*. In effect, the models test the null hypothesis that, all else equal, a male should prefer a roving male strategy whenever this maximises his access to females and the siring rate he can achieve, and it does so at the level of the population rather than the species. As with all such exercises, we seek to establish the circumstances under which all else is not equal. The difference in the payoffs for the two options provides us with a direct measure of the relative magnitude of the selection pressure favouring whichever strategy males actually pursue, and hence of the male’s “regret” (the opportunity cost he bears by not pursuing the all-else-equal optimal strategy). This, in turn, gives us insight into the magnitude of the advantages that being social (monogamous, in this case) must provide.

The present analyses indicate that monogamous species do not consistently occupy territories that are absolutely larger than those defended by polygamous species. In fact, with the possible exception of *Pithecia* and *Lepilemur*, the females of monogamous species occupy much smaller territories than those defended by the males of polygamous species (Fig. 1a). Even on a per capita metabolic basis, it is only some callitrichid genera (with *Callithrix* as an exception) that have *relatively* larger territories than the polygamous genera; for all the other monogamous genera, metabolic-adjusted territory sizes are indistinguishable from those of polygamous primates (Fig. 1b). Given the mobility criterion that defines territoriality in primates, males of these species could easily defend territories large enough to include the ranges of 5-7 solitary females (Figs. 2 and 4). Even if females are only briefly in oestrus and males risk losing some of them to rivals, they would, with the possible exception of *Nomascus* and *Leontopithecus*, all benefit by pursuing a roving strategy (Table 1). *Nomascus* and *Leontopithecus* occupy more marginal habitats compared to other members of their respective families (Dunbar et al. 2019; Rylands 1989; Kierulff & Rylands 2003). Given these results and the fact that evidence from models of time budgets (Dunbar et al. 2009, 2019; Korstjens et al. 2018) suggests that, except on the limits of their biogeographical distributions, most primates are not under as much ecological pressure as has usually been assumed, this effectively rules out any likelihood that the females of monogamous species are forced to live alone due to ecological competition.

Brotherton & Manser (1997) suggested that the problem might be that males are unable to prevent rivals mating with their females if several are in oestrus simultaneously in different locations. That this might be an issue is suggested by *Cercopithecus mitis*. Although their groups conventionally have a single breeding male, they are subject to periodic invasions by bands of roving males when several females are in oestrus simultaneously (Henzi & Lawes 1987; Cords 2004; Roberts et al. 2014; Gao & Cords 2020); these roving males can gain up to 40% of sirings (Roberts et al. 2014). Guenon groups typically contain ~7 females, compared to the 3-4 characteristic of the unimale groups of *Colobus* and *Gorilla* (Dunbar et al. 2018b) and lie on the cusp of where a single male can successfully defend a group of females (Andelman 1987). So long as there is only one female in oestrus, the male can defend his group against rivals; but when several females’ cycles coincide, he is unable to prevent other males gaining access to some of the females (Dunbar 2000, 2019; see also Pawlowski et al. 1998).

It is important to remember that current circumstances are not always relevant to the evolutionary origins of a trait, since they often involve subsequent accommodations (through the exploitation of windows of evolutionary opportunity) after the trait has evolved. The callitrichids provide a case in point. Analysis of the possible routes to monogamy with biparental care in these taxa suggests that biparental care evolved *after* pairbonding, and that pairbonding in turn allowed the females to evolve twinning with much shorter interbirth intervals (Dunbar 1995a). Once the female can produce twins twice a year, the male is effectively locked into monogamy because it means he has to be able to control matings to more than four females to make roving worth his while. There is, thus, an important distinction between whether roving is worth a male’s while under *current* circumstances when females can twin twice a year and whether it was worthwhile under the *ancestral* conditions when females had a conventional primate lifehistory and produced only singleton infants on the same schedule as the smaller platyrrhines. Reanalysis using a single female as the baseline suggests that, under the ancestral conditions, the callitrichids’ payoff ratio for roving does not differ from those for the other hylobatid and cebid genera: roving is consistently advantageous (Table 1; Fig. S7). Notice, by the way, that roving is always advantageous for *Saguinus* but at best only marginally so for *Callithrix* (Table 1). This is in line with the empirical evidence that *Callithrix* groups are much more stable than those of *Saguinus*, whose males often adopt what is, in effect, a roving strategy in which breeding males sometimes move on to another group after the female has conceived, leaving the helpers-at-the-nest to rear their offspring and inherit the female (Goldizen 1988, 1990; Ferrari & Digby 1996).

In sum, while solitary foraging by females is clearly a necessary condition for the evolution of monogamy, it is not a sufficient explanation. The substantive issue, then, is why the males opt for monogamy when they would do much better by roving – especially so given that prosimian males do exactly this. The suggestion that monogamous males are protecting mating access to the female is implausible, given interbirth intervals that can be as long as 39 months in the case of gibbons and the magnitude of the fitness differentials between monogamous and roving males (which can be as much as 10-fold: Figs. 2 and 4). If breeding is highly synchronised, males might be forced into monogamy simply because they cannot both search and keep rivals away from two females simultaneously (Knowlton 1979). However, comparison of Figs. S5 and S6 suggests that this only works if there is a breeding season. Although most of the genera in this sample breed throughout the year (*Colobus*: Fashing 2007; *Cercopithecus*: Isbell et al. 2004, except perhaps *C. mitis*; gibbons: Leighton 1987; Savini et al. 2008; *Gorilla*: Stewart & Harcourt 1987), many of the platyrrhine genera (callitrichids: Dietz & Baker 1993; Digby et al. 2007; *Aotus*: Fernandez-Duque 2007; pithecids: Norcronk 2007) and the klipspringer (Norton 1980) have birth (and therefore mating) seasons. Nonetheless, this would make a social strategy worthwhile only in the case of the callitrichids, and then only because the females have evolved the capacity to twin up to twice a year. Once again, this suggests that callitrichid females have endeavoured to lock their males into stable monogamy (Dunbar 1995a). For the other genera, the fact that primates, in particular, have a long period of offspring dependency suggests that the problem is more likely to be associated with offspring survival.

The models provide us with an explicit estimate of the opportunity cost that males pay by being obligate monogamists. The results suggest that this cost is extremely high (potentially as high as 4-10 offspring per reproductive cycle). In contrast, the nocturnal prosimians are willing to opt for roving male promiscuity even though the benefits of doing so are quite modest (on average, just 0.72 extra offspring per reproductive cycle: Fig. 2). In addition to this opportunity cost, monogamous mammals pay an additional energetic cost. Across mammals and birds, as well as primates, brain size is significantly larger in monogamous species than it is in promiscuous species (Shultz & Dunbar 2007; Dunbar 2010), probably because of the cognitive demands of maintaining stable relationships (see Dunbar 2018; Dunbar & Shultz 2021). These species thus bear a double cost: they sacrifice sirings and they incur an expensive neuro-cognitive cost that asocial roving males like the prosimians avoid.

These costs equate to the selection in favour of monogamy, and their magnitude suggests that the fitness advantages of monogamy must be very substantial. What those advantages might be have not been explored here. The only two costs known to be high for anthropoids that might function as a countervailing benefit are predation risk (Cheney & Wrangham 1987; Isbell 1994; Shultz et al. 2004; Shultz & Finlayson 2010; see also Burnham et al. 2012) and infanticide risk (van Schaik & Dunbar 1990; van Schaik & Kappeler 1997; Borries et al. 2011; Opie et al. 2013; Lukas & Huchard 2014; Lowe et al. 2019). Predation risk must play some role, since it is the principal driver for group-living in primates (Shultz et al. 2004; Shultz & Finlayson 2010). However, the fact that the females of monogamous genera are willing to forage alone suggests that the risk of predation is likely to be low.

Primates, in contrast to most other mammals, run a high risk of infanticide because of their greatly extended interbirth intervals, in turn a direct consequence of their unusually large brains. Infanticide is significantly lower in monogamous primates than for primates that live in polygamous groups (Borries et al. 2011; Opie et al. 2013; Lukas & Huchard 2014), strongly suggesting there has been selection to minimise this risk by adopting monogamy. That infanticide is a significant risk at least in gibbons is demonstrated by Ma et al. (2019): in a wide-ranging review, they found that 50% of gibbon infants died or disappeared within two months of the pair male being replaced. On top of natural mortality, this represents a very substantial fitness cost to a male who may only sire around five offspring in a reproductive career.

## Online Supplementary Material

### Comparison of Mitani-Rodman and Lowen-Dunbar indices of territoriality

The Mitani-Rodman territorial defendability index (Eq. [1]) and the equivalent Lowen-Dunbar index (Eq. [3]) are based on slightly different assumptions about how a male detects intruders into his territory. The first assumes that the male searches throughout his territory for intruders and indexes the male’s ability to defend a territory of a given size in terms of the frequency with which he can visit all parts of his territory while foraging randomly; the second assumes that males detect intruders mainly along their boundary before they enter the body of the territory itself, and so indexes the frequency with which the male can visit all segments of the boundary. For mathematical convenience, both assume that territories are perfect circles. In both cases, the critical value of the index that best differentiates territorial from non-territorial species is determined empirically.

Figure S1 plots the maximum number of defendable females predicted by the Lowen-Dunbar inequality (Eq. [4]) against the equivalent value predicted by the Mitani-Rodman inequality (Eq. [2]), for detection arcs on the territory boundary of (a) s=0.05 km, (b) s=0.2 km and (c) s=0.5 km. Relative to the Mitani-Rodman index, the Lowen-Dunbar index underestimates the defendable area (and hence the maximum number of defendable females) when the detection distance is very short (*s*=0.05 km, or 25 m either side of the male) and overestimates it when the detection distance is long (*s*=0.5 km, or 250 m either side of the male). The best fit is provided by a detection arc in the order of *s*=0.2 km (i.e. 100 m either side of the male).

**Figure S1.**
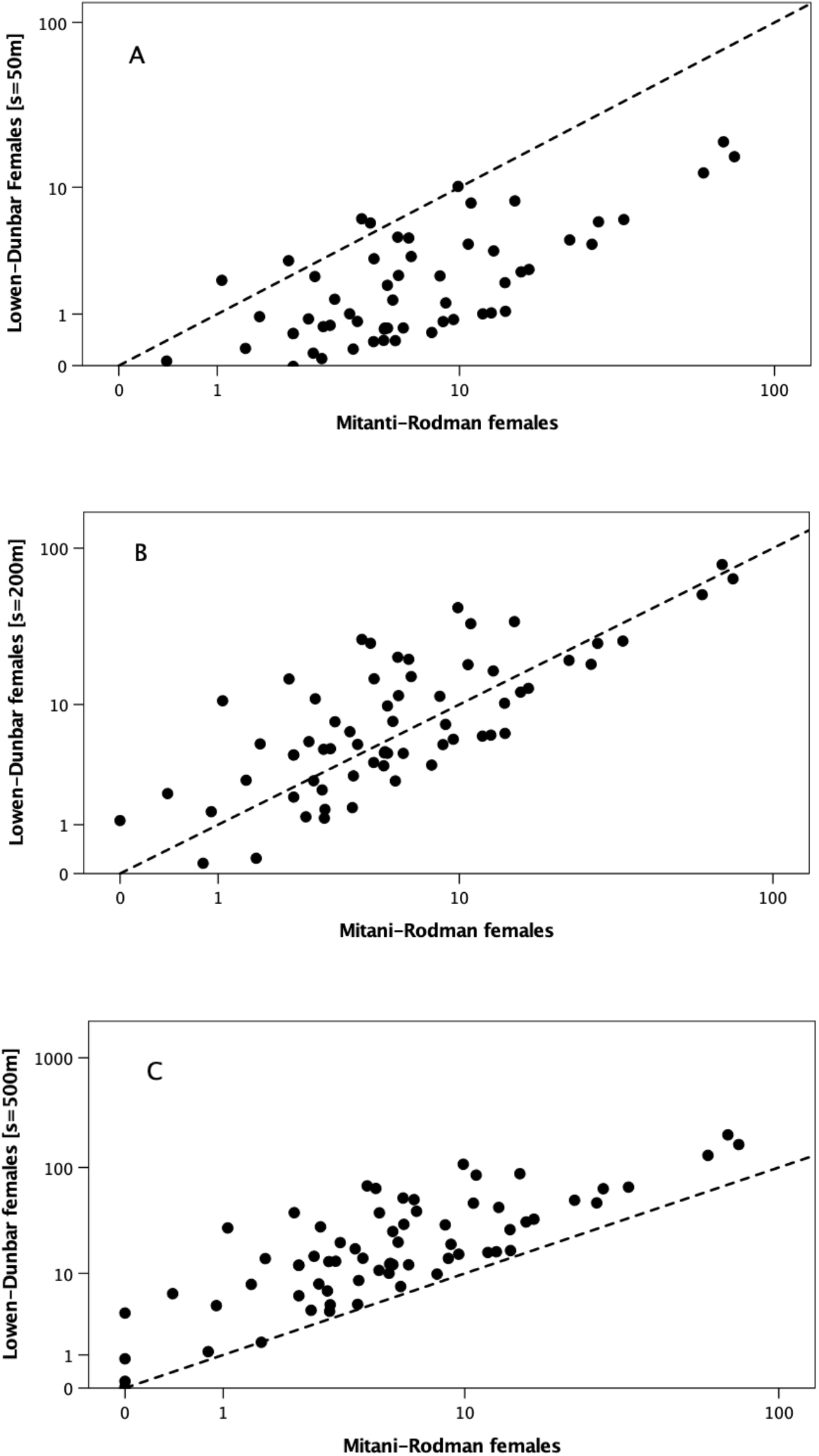
Comparison of the maximum number of defendable females predicted by the two defendability indices, with detection arc in the Lowen-Dunbar equation set at (a) s= 0.05 km, (b) s=0.2 km and (c) s=0.5 km. Dashed line is the line of equality.

### Lowen-Dunbar index prediction of number of defendable females

Figure S2 plots the ratio of the maximum number of defendable females predicted by the Lowen-Dunbar index of defendability relative to the observed number of females in the group (with appropriately adjusted values for the callitrichids). When ratio≤1, males should prefer to be social (i.e. permanently attached to a female or group of females), and when ratio>1 they should prefer roving male polygamy. *Nomascus, Leontopithecus*, *Saguinus* and all the group-living polygamous genera should prefer to be social. The other monogamous species, including the antelope *Oreotragus* should prefer to rove.

**Figure S2.**
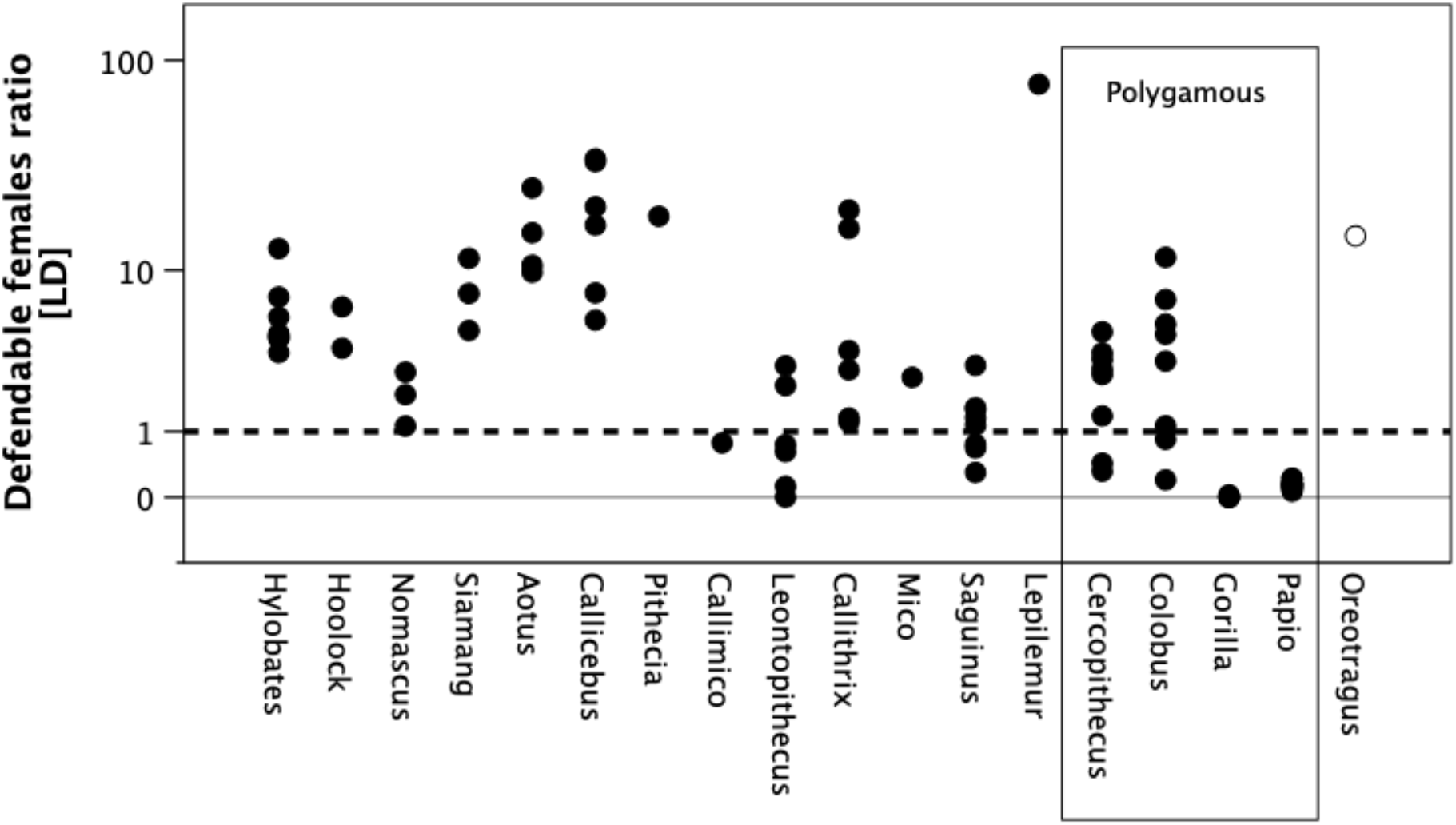
Maximum number of defendable females that a territorial male can have, given the length of his day journey, estimated by the Lowen-Dunbar index with s=200 m, as a ratio of the observed female group size for the population. Unfilled symbol: ungulate.

### Effect of detection distance on rover payoff ratio

Figure 4 assumes that a male can detect a female who is within a 200m-wide path as he forages. Figure S3 plots the payoff ratios for the various populations when the detection range is either 2*r=*50 m or 2*r*=500 m. One-sample t-tests testing the null hypothesis that *E*(*p*)=1 are given in Table S1. The distribution of datapoints remains similar, but payoff ratios are inevitably less advantageous to rovers when the detection distance is shorter. Even so, roving is still advantageous for most genera. When the detection range is as large as 2*r*=500 m (realistically, only possible in open terrestrial habitats), roving is advantageous for all genera except *Gorilla.* At these distances, it would even pay *Colobus* and *Papio* males to adopt a roving strategy.

**Figure S3.**
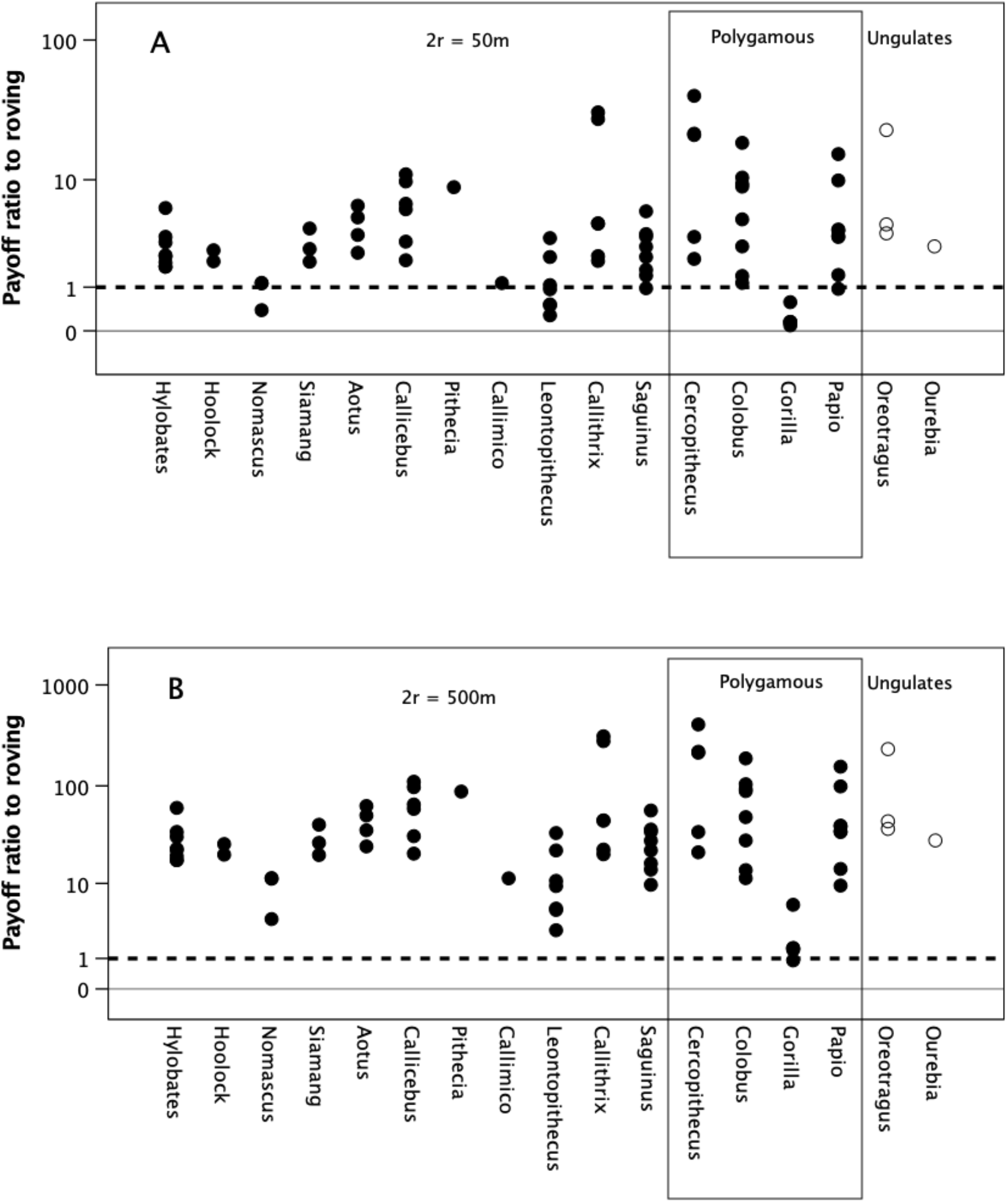
Payoff ratio favouring roving predicted by Eq. [8] with the detection distance set to *(a)* 2r = 50 m and (b) 2*r* = 500 m. When the payoff ratio >1, males should prefer a roving strategy.

**Table S1.**
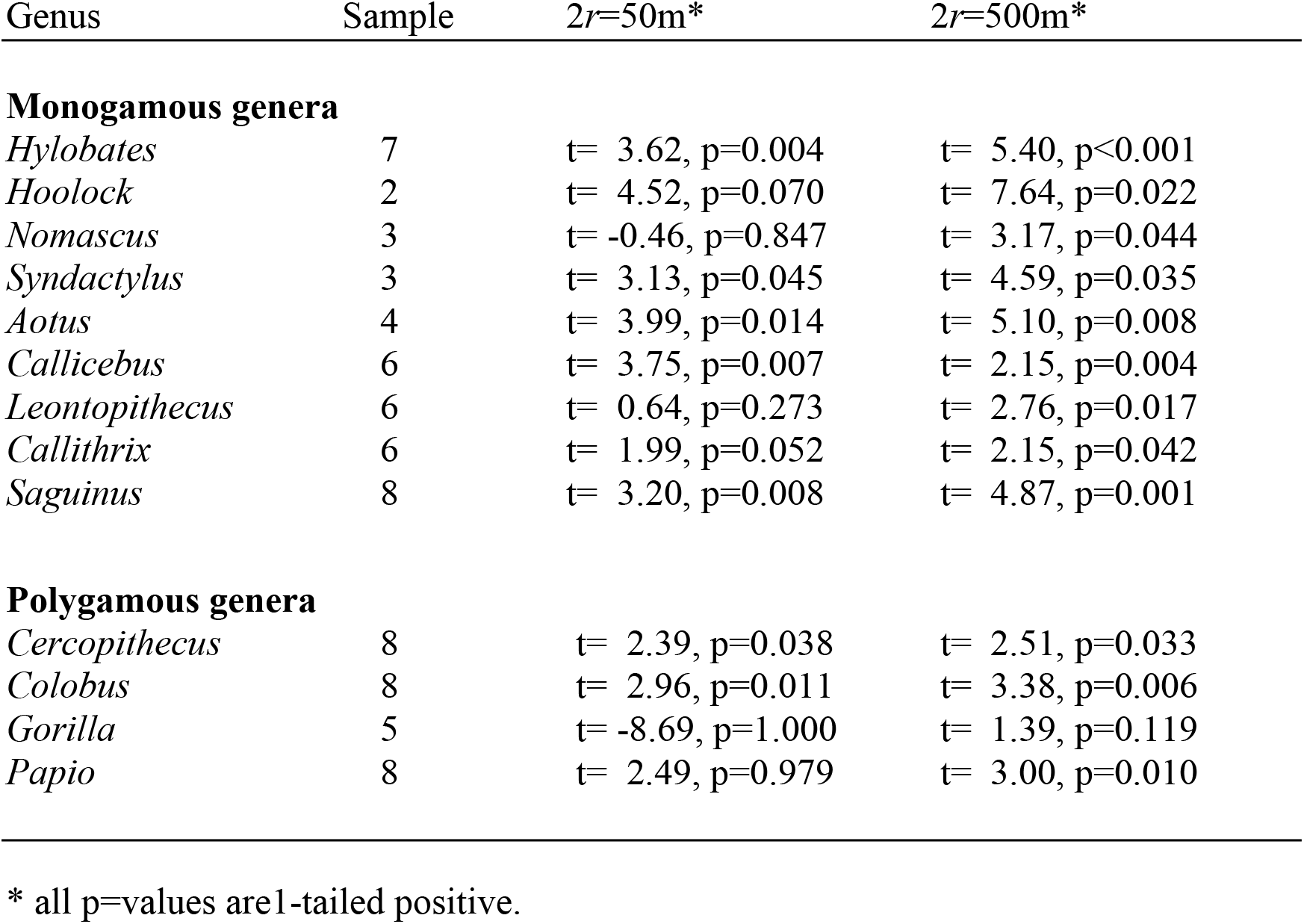
One-sample t-tests for deviation from *E*(*p*)=1.

### Effect of breeding synchrony on payoff ratios

Figure S4 plots the distribution of the proportion of co-cycling females across the genera, as determined by Eq. [8]. Female reproductive cycles are assumed to be unsynchronised. Most genera have a very low likelihood that two or more females would be in oestrus at the same time. This is mainly because the number of females involved is small relative to the length of the interbirth interval. Figure S5 plots the payoff ratios when these are adjusted for the number of co-cycling females (assuming that the male can monopolise only one of these at any one time). The differences compared to Figure 4 are marginal: across all 71 populations, the mean loss is approximately one siring per reproductive cycle (means of 10.73±13.9 vs 9.85±11.5 sirings).

**Figure S4.**
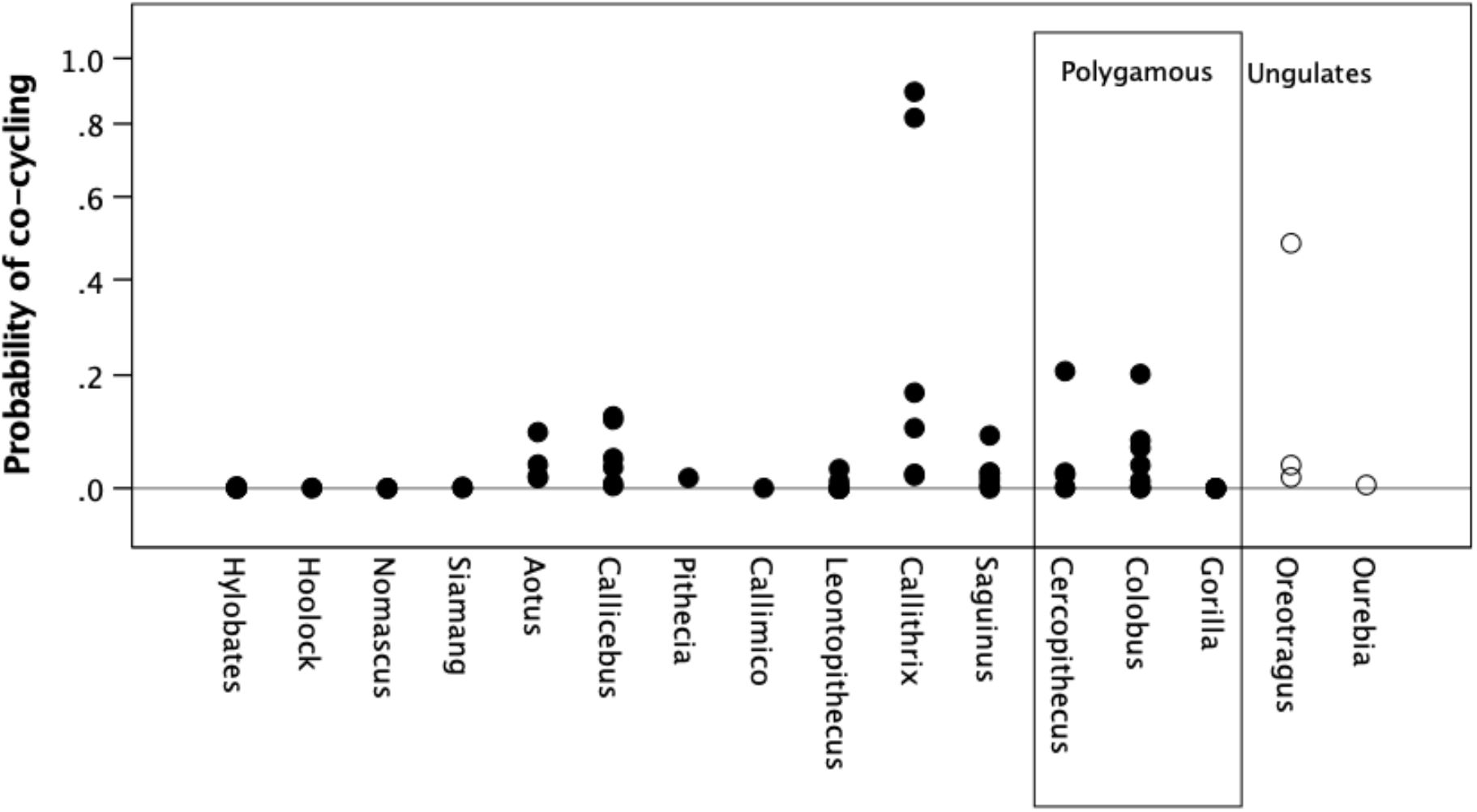
Summed probability that two or more of the females that a male locates during the course of the average interbirth interval will be in oestrus at the same time.

**Figure S5.**
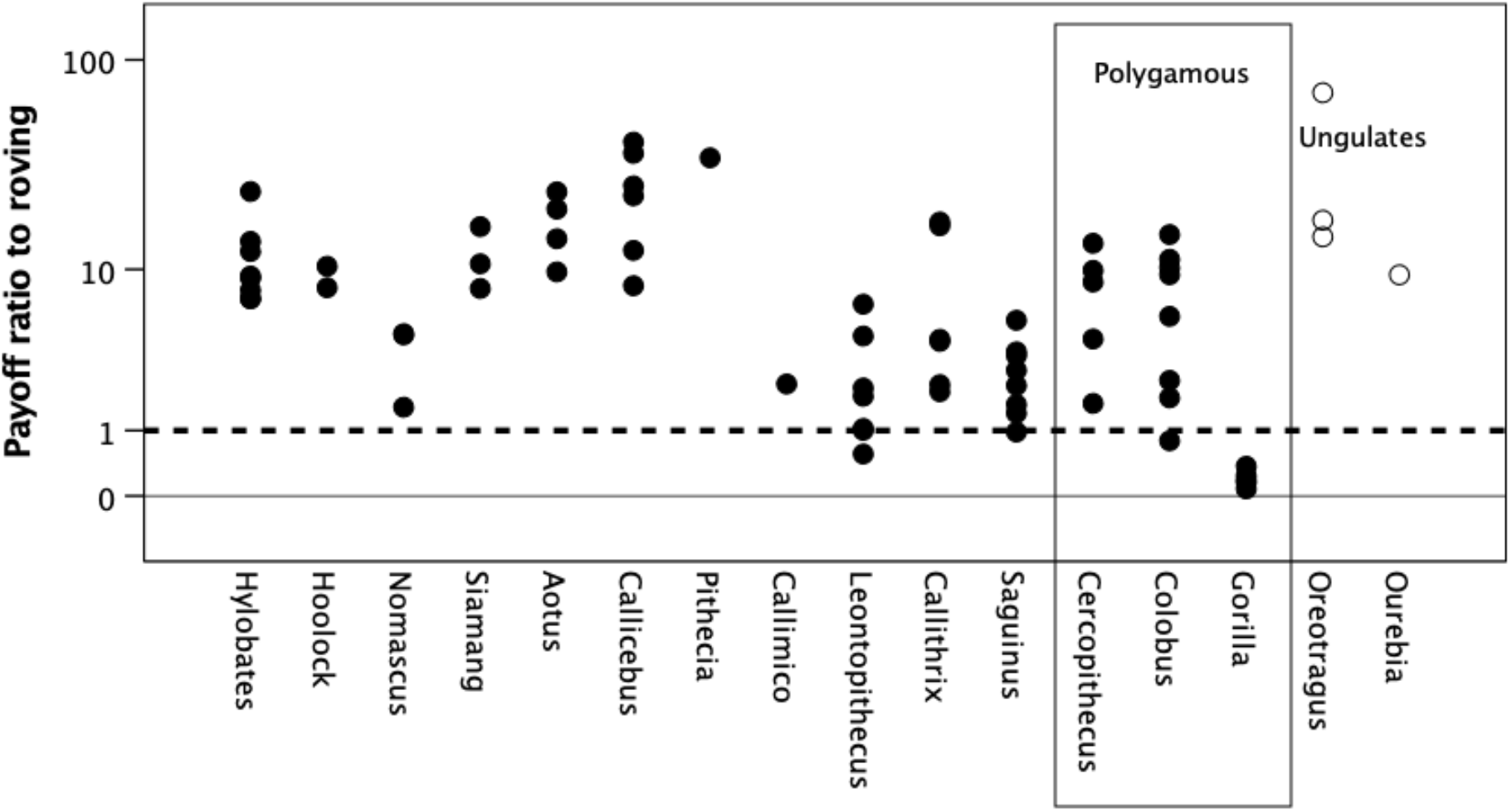
Payoff ratio to rovers adjusted for the number of females who would be oestrus at the same time, such that the male could not monopolise both (calculated from Eq. [7]). Timebase is population interbirth interval. Unfilled symbols: ungulates.

Fig. S6 plots the effect that a six-month breeding season would have on the payoff ratio when 2*r*=200m, adjusted for the probability that some of the females will be co-cycling. A 6-month birth season reduces the mean payoff ratio by approximately one further siring compared to Fig. S5 (means of 9.85±11.5 vs 8.95±9.6 sirings).

**Figure S6.**
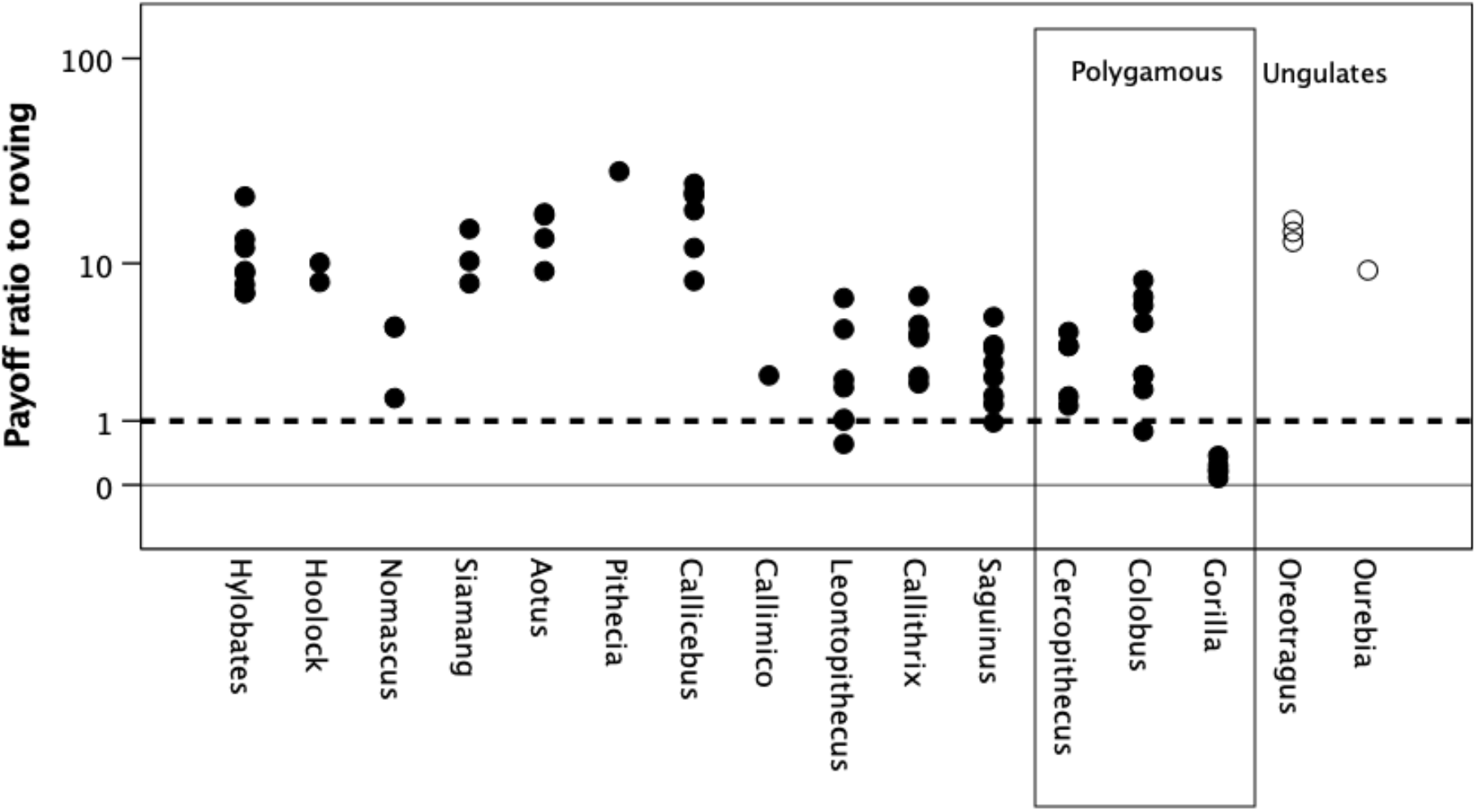
Payoff ratios to roving adjusted for co-cycling (as for Figure S6) and for a 6-month mating season. Unfilled symbols: ungulates.

### Setting baseline to observed female group size for Callitrichids

Figure 4 considers the current circumstances in which callitrichid females are able to twin up to twice a year. In effect, this considers the situation in which females have evolved further strategies that lock the male into biparental care. Figure S7 plots the same payoff ratios as in Figure 4, but adds (as unfilled symbols) the number of defendable females that callitrichid males would gain under the presumed ancestral condition in which females produced singleton births once a year (the ancestral anthropoid condition). When the baseline is one female, all the callitrichid genera would benefit by roving, and, as a group, do not differ from any of the hylobatids and cebids.

**Figure S7.**
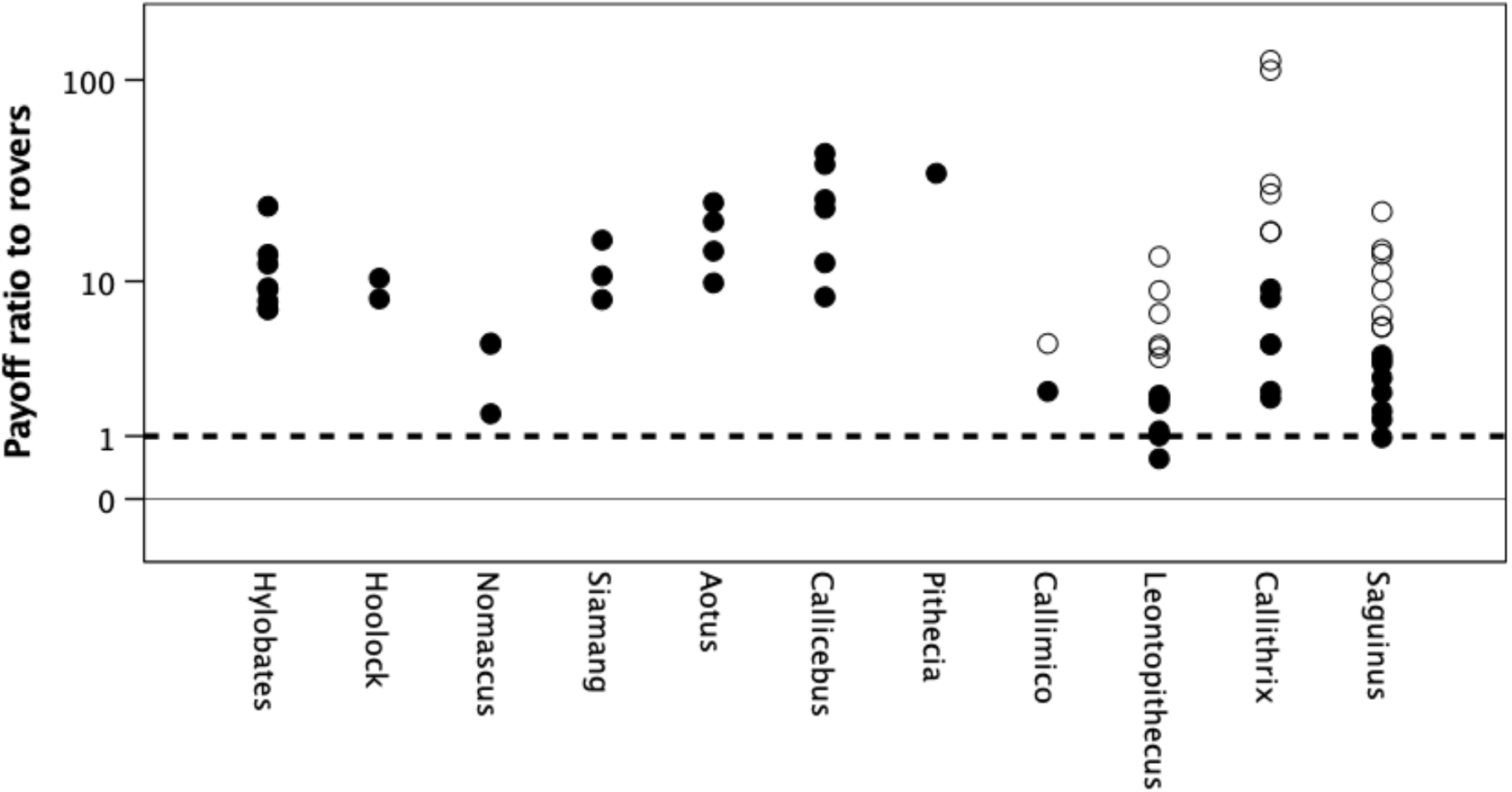
Payoff ratio to rovers when the detection distance 2r=200 m. Filled symbols are compared to the same baseline as Figure 4 (2 or 4 female-equivalents for the callitrichids); unfilled symbols compare the rover’s payoff for the callitrichid genera, assuming that in the ancestral condition females would only have been able to produce singleton young once a year.

## References

Abbott, D. H. (1984). Behavioral and physiological suppression of fertility in subordinate marmoset monkeys. American Journal of Primatology 6: 169–186.

Adamczak, V. & Dunbar, R.I.M. (2008). Variation in the mating system of oribis and their ecological determinants. African Journal of Ecology, 45, 197–206.

Andelman, S. (1987). Ecological and social determinants of cercopithecine mating patterns. In: D.I. Rubenstein & R.W. Wrangham (eds.) Ecological Aspects of Social Evolution, pp. 201–217. Princeton, NJ: Princeton University Press.

Bales, K., O’Herron, M., Baker, A. J., & Dietz, J. M. (2001). Sources of variability in numbers of live births in wild golden lion tamarins (Leontopithecus rosalia). American Journal of Primatology: Official Journal of the American Society of Primatologists, 54(4), 211–221.

Bettridge, C., Lehmann, J. & Dunbar, R.I.M. (2010). Trade-offs between time, predation risk and life history, and their implications for biogeography: a systems modelling approach with a primate case study. Ecological Modelling 221: 777–790.

Borries, C., Savini, T. & Koenig, A. (2011). Social monogamy and the threat of infanticide in larger mammals. Behavioral Ecology and Sociobiology, 65, 685–693.

Brotherton, P. N.M. & Komers, P.E. (2003). Mate guarding and the evolution of social monogamy in mammals. In: U.H. Reichard & C. Boesch (eds.) Monogamy: Mating Strategies and Partnerships in Birds, Humans and Other Mammals, pp 42–58. Cambridge: Cambridge University Press.

Brotherton, P.N. & Manser, M.B. (1997). Female dispersion and the evolution of monogamy in the dik-dik. Animal Behaviour 54: 1413–1424.

Burnham, D., Bearder, S., Cheyne, S., Dunbar, R.I.M., Macdonald, D. (2012). A taste of predation past: is nocturnal behaviour in primates explained by the threat of predation by cats? Folia Primatologica, 83, 236–251.

Campbell, C.J., Fuentes, A., Mackinnon, K., Panger, M. & Bearder, S.K. (eds.) (2007). Primates in Perspective. Oxford: Oxford University Press.

Cheney, D.L. & Wrangham, R.W. (1987). Predation. In: B. Smuts, D. Cheney, R. Seyfarth, R.W. Wrangham & T. Struhsaker (eds.), Primate Societies, pp. 227–239. Chicago: Chicago University Press.

Cords, M. (2004). When are there influxes in blue monkey groups?. In The Guenons: Diversity and Adaptation in African Monkeys, pp. 189–201. Springer, Boston MA.

Davies, N.B. (1992). Dunnock Behaviour and Social Evolution. Oxford: Oxford University Press.

Dietz, J. M., & Baker, A. J. (1993). Polygyny and female reproductive success in golden lion tamarins, *Leontopithecus rosalia*. Animal Behaviour, 46, 1067–1078.

Dietz, J. M., Baker, A. J., & Miglioretti, D. (1994). Seasonal variation in reproduction, juvenile growth, and adult body mass in golden lion tamarins (*Leontopithecus rosalia*). American Journal of Primatology, 34, 115–132.

Digby, L.J., Ferrari, S.F. & Saltzman, W. (2007). The role of competition in cooperatively breeding species. In: C. Campbell, A. Fuentes, K. Mackinnon, S.K. Bearder & S.K. Stumpf (eds.) Primates in Perspective, pp. 85–106. New York: Oxford University Press.

Dobson, F. S., Way, B. M., & Baudoin, C. (2010). Spatial dynamics and the evolution of social monogamy in mammals. Behavioral Ecology, 21(4), 747–752.

Doran, D.M. & McNeilage, A. (1998). Gorilla ecology and behaviour. Evolutionary Anthropology, 6, 120–131.

Doran-Sheehy, D.M. & Boesch, C. (2004). Behavioral ecology of western gorillas: new insights from the field. American Journal of Primatology, 64, 139–143.

Dunbar, R.I.M. (1988). Primate Social Systems. London: Chapman & Hall.

Dunbar, R.I.M. (1992). Time: a hidden constraint on the behavioural ecology of baboons. Behav. Ecol. Sociobiol. 31: 35–49.

Dunbar, R.I.M. (1995a). The mating system of Callitrichid primates. I. Conditions for the coevolution of pairbonding and twinning. Animal Behaviour, 50, 1057–1070.

Dunbar, R.I.M. (1995b). The mating system of Callitrichid primates. II. The impact of helpers. Animal Behaviour, 50, 1071–1089.

Dunbar, R.I.M. (2000). Male mating strategies: a modelling approach. In: P. Kappeler (ed.) Primate Males, pp. 259–268. Cambridge: Cambridge University Press.

Dunbar, R.I.M. (2001). The economics of male mating strategies among primates. In: J. van Hooff, R. Noë & P. Hammerstein (eds.) Economic Models of Animal and Human Behaviour, pp. 245–269. Cambridge: Cambridge University Press.

Dunbar, R.I.M. (2010). Deacon’s dilemma: the problem of pairbonding in human evolution. In: R.I.M. Dunbar, C. Gamble & J.A.J. Gowlett (eds.) Social Brain, Distributed Mind, pp. 159–179. Oxford: Oxford University Press.

Dunbar, R.I.M. (2018). The anatomy of friendship. Trends in Cognitive Sciences, 22, 32–51.

Dunbar, R.I.M. (2019). Fertility as a constraint on group size in African great apes. Biological Journal of the Linnaean Society 129: 1–13.

Dunbar, R.I.M. & Dunbar, P. (1980). The pairbond in klipspringer. Animal Behaviour, 28, 251–263.

Dunbar, R.I.M. & Shultz, S. (2010). Bondedness and sociality. Behaviour 147: 775–803.

Dunbar, R.I.M. & Shultz, S. (2021). Social complexity and the fractal structure of social groups in primate social evolution. Biology Reviews.

Dunbar, R.I.M., Buckland, D. & Miller, D. (1990). Mating strategies of male feral goats: a problem in optimal foraging. Animal Behaviour, 40, 653–667.

Dunbar, R.I.M., Korstjens, A.H. & Lehmann, J. (2009). Time as an ecological constraint. Biological Reviews, 84, 413–429.

Dunbar, R.I.M., MacCarron, P. & Robertson, C. (2018a). Tradeoff between fertility and predation risk drives a geometric sequence in the pattern of group sizes in baboons. Biology Letters 14: 20170700.

Dunbar, R.I.M., MacCarron, P. & Shultz, S. (2018b). Primate social group sizes exhibit a regular scaling pattern with natural attractors. Biology Letters 14: 20170490.

Dunbar, R.I.M., Cheyne, S., Lan, D., Korstjens, A.H., Lehmann, J. & Cowlishaw, G. (2019). Environment and time as constraints on the biogeographical distribution of gibbons. American Journal of Primatology, 81, e22940.

Emlen, S. T., & Oring, L. W. (1977). Ecology, sexual selection, and the evolution of mating systems. Science 197: 215–223.

Enstam, K.L. & Isbell, L.A. (2007). The guenons (genus *Cercopithecus*) and their allies: behavioural ecology of polyspecific associations. In: C.J. Campbell, A. Fuentes, K. Mackinnon, M. Panger & S.K. Bearder (eds.) Primates in Perspective, pp. 252–273. Oxford: Oxford University Press.

Fashing, P.J. (2007). African colobine monkeys: patterns of between-group interaction. C.J. Campbell, A. Fuentes, K. Mackinnon, M. Panger & S.K. Bearder (eds.) Primates in Perspective, pp. 201–223. Oxford: Oxford University Press.

Fernandez-Duque, E. (2007). Aotinae: social monogamy in the only nocturnal haplorhines. In: C.J. Campbell, A. Fuentes, K. Mackinnon, M. Panger & S.K. Bearder (eds.) Primates in Perspective, pp. 139–154. Oxford: Oxford University Press.

Ferrari, S. F., & Digby, L. J. (1996). Wild Callithrix groups: stable extended families?. American Journal of Primatology, 38(1), 19–27.

Gao, L., & Cords, M. (2020). Effects of female group size on the number of males in blue monkey (*Cercopithecus mitis*) groups. International Journal of Primatology 00: 1–18.

Goldizen, A.W. (1988). Tamarin and marmoset mating systems: unusual flexibility. Trends in Ecology and Evolution, 3, 36–40.

Goldizen, A. W. 1990. A comparative perspective on the evolution of tamarin and marmoset social systems. International Journal of Primatology 11: 63–83.

Hamilton, W. D. (1964). The genetical evolution of social behaviour. I, II. Journal of Theoretical Biology, 7: 1–52.

Harvey, P.H. & Clutton-Brock, T.H. (1985). Life history variation in primates. Evolution, 39, 559–581.

Henzi, S.P. & Lawes, M. (1987). Breeding season influxes and the behaviour of adult male samango monkeys (*Cercopithecus mitis albogularis*). Folia Primatologica, 48, 125–136.

Hill, R.A., Lycett, J., & Dunbar, R.I.M. (2000). Ecological determinants of birth intervals in baboons. Behavioral Ecology, 11, 560–564.

Hilgartner, R., Fichtel, C., Kappeler, P. M., & Zinner, D. (2012). Determinants of pair-living in red-tailed sportive lemurs (Lepilemur ruficaudatus). Ethology, 118(5), 466–479.

Isbell LA (1994). Predation on primates: ecological patterns and evolutionary consequences. Evolutionary Anthropology 3: 61–71.

Isbell, L.A., Cheney, D.L. & Seyfarth, R.M. (2004). Why vervet monkeys (*Cercopithecus aethiops*) live in multimale groups. In: The Guenons: Diversity and Adaptation in African Monkeys (pp. 173–187). Boston, MA: Springer.

Kamilar, J.M. & Cooper, N. (2013). Phylogenetic signal in primate behaviour, ecology and life history. Philosophical Transactions of the Royal Society, London, 368B, 20120341.

Kappeler, P. M., & Pozzi, L. (2019). Evolutionary transitions toward pair living in nonhuman primates as stepping stones toward more complex societies. Science advances, 5(12), eaay1276.

Kierulff, M. C. M., & Rylands, A. B. (2003). Census and distribution of the golden lion tamarin (*Leontopithecus rosalia*). American Journal of Primatology 59, 29–44.

Knowlton, N. (1979). Reproductive synchrony, parental investment, and the evolutionary dynamics of sexual selection. Animal Behaviour, 27, 1022–1033.

Komers, P.E. & Brotherton, P.N.M. (1997). Female space use is the best predictor of monogamy in mammals. Proceedings of the Royal Society, London, 264B, 1261–1270.

Korstjens, A.H. & Dunbar, R.I.M. (2007). Time constraints limit group sizes and distribution in red and black-and-white colobus monkeys. Int. J. Primatol. 28: 551–575.

Korstjens, A.H., Lehmann, J. & Dunbar, R.I.M. (2018). Time constraints do not limit group size in arboreal guenons but do explain community size and distribution patterns. International Journal of Primatology, 39, 511–531.

Krause, J. & Ruxton, G.D. (2000). Living in Groups. Oxford: oxford university Press.

Lehmann., J., Korstjens, A.H. & Dunbar, R.I.M. (2008a). Time management in great apes: implications for gorilla biogeography. Evolutionary Ecology Research 10: 515–536.

Leighton, D.M. (1987). Gibbons: territoriality and monogamy. In: Smuts, B.B., D.L. Cheney, R.M. Seyfarth, R.W. Wrangham & T.T. Strusaker (eds.) Primate Societies, pp.135–145. Chicago: Chicago University Press.

Ligon, J.D. (1999). The Evolution of Avian Breeding Systems. Oxford: Oxford University Press

Lowe, A.E., Hobaiter, C., Asiimwe, C., Zuberbühler, K., & Newton-Fisher, N.E. (2019). Intra-community infanticide in wild, eastern chimpanzees: a 24-year review. Primates (in press).

Lowen, C.B. & Dunbar, R.I.M. (1994). Territory size and defendability in primates. Behavioral Ecology and Sociobiology, 35, 347–354.

Lukas, D. & Clutton-Brock, T.H. (2013). The evolution of social monogamy in mammals. Science, 341, 526–530.

Lukas, D. & Huchard, E. (2014). The evolution of infanticide by males in mammalian societies. Science, 346, 841–844.

Ma, C.Y., Brockelman, W.Y., Light, L.E., Bartlett, T.Q., & Fan, P.F. (2019). Infant loss during and after male replacement in gibbons. American Journal of Primatology, 81, e23036.

Mitani, J.C. & Rodman, P. (1979). Territoriality: the relation of ranging pattern and home range size to defendability, with an analysis of territoriality among primate species: Behavioral Ecology and Sociobiology, 5, 241–251.

Müller, A. E., & Thalmann, U. R. S. (2000). Origin and evolution of primate social organisation: a reconstruction. Biological Reviews, 75(3), 405–435.

Norconk, M.A. (2007). Sakis, uakaris, and titi monkeys: behavioral diversity in a radiation of primate seed predators. In: C.J. Campbell, A. Fuentes, K. Mackinnon, M. Panger & S.K. Bearder (eds.) Primates in Perspective, pp. 123–138. Oxford: Oxford University Press.

Opie, C., Atkinson, Q., Dunbar, R.I.M. & Shultz, S. (2013). Male infanticide leads to social monogamy in primates. PNAS, 110, 13328–13332.

Overdorff, D. J., & Tecot, S. R. (2006). Social pair-bonding and resource defense in wild red-bellied lemurs (Eulemur rubriventer). In Lemurs (pp. 235–254). Springer, Boston, MA.

Pawlowski, B.P., Lowen, C.B. & Dunbar, R.I.M. (1998). Neocortex size, social skills and mating success in primates. Behaviour, 135, 357–368.

Porter, L. M., Hanson, A. M., & Becerra, E. N. (2001). Group demographics and dispersal in a wild group of Goeldi’s monkeys (Callimico goeldii). Folia Primatologica, 72(2), 108–110.

Robbins, M. (2007). Gorillas: diversity in ecology and behaviour. In: C.J. Campbell, A. Fuentes, K. Mackinnon, M. Panger & S.K. Bearder (eds.) Primates in Perspective, pp. 305–321. Oxford: Oxford University Press.

Roberts, S.J., Nikitopoulos, E. & Cords, M. (2014). Factors affecting low resident male siring success in one-male groups of blue monkeys. Behavioral Ecology, 25, 852–861.

Rutberg, A. T. (1983). The evolution of monogamy in primates. Journal of Theoretical Biology, 104(1), 93–112.

Rylands, A. B. (1989). Sympatric Brazilian callitrichids: the black tufted-ear marmoset, *Callithrix kuhli*, and the golden-headed lion tamarin, *Leontopithecus chrysomelas*. Journal of Human evolution, 18, 679–695.

Savini, T., Boesch, C., & Reichard, U. H. (2008). Home-range characteristics and the influence of seasonality on female reproduction in white-handed gibbons (*Hylobates lar*) at Khao Yai National Park, Thailand. American Journal of Physical Anthropology, 135, 1–12.

van Schaik, C. & Dunbar, R.I.M. (1990). The evolution of monogamy in large primates: a new hypothesis and some critical tests. Behaviour, 115, 30–62.

van Schaik, C.P. & Kappeler, P.M. (1997). Infanticide risk and the evolution of male–female association in primates. Proceedings of the Royal Society, London, 264B, 1687–1694.

Shultz, S. & Dunbar, R.I.M. (2007). The evolution of the social brain: Anthropoid primates contrast with other vertebrates. Proceedings of the Royal Society, London, 274B: 2429–2436.

Shultz, S. & Dunbar, R.I.M. (2010). Social bonds in birds are associated with brain size and contingent on the correlated evolution of life-history and increased parental investment. Biological Journal of the Linnaean Society 100:111–123.

Shultz, S. & Finlayson, L.V. (2010). Large body and small brain and group sizes are associated with predator preferences for mammalian prey. Behavioral Ecology, 21, 1073–1079.

Shultz, S., Opie, C. & Atkinson, Q.D. (2011). Stepwise evolution of stable sociality in primates. Nature, 479, 219–222.

Shultz, S., Noe, R., McGraw, S. & Dunbar, R.I.M. (2004). A community-level evaluation of the impact of prey behavioural and ecological characteristics on predator diet composition. Proceedings of the Royal Society, London, 271B, 725–732.

Stewart, K.J. & Harcourt, A.H. (1987). Gorillas: variation in female relationships. In: Smuts, B.B., D.L. Cheney, R.M. Seyfarth, R.W. Wrangham & T.T. Strusaker (eds.) Primate Societies, pp.155–164. Chicago: Chicago University Press.

Williams, J. M., Oehlert, G. W., Carlis, J. V., & Pusey, A. E. (2004). Why do male chimpanzees defend a group range? Animal Behaviour, 68(3), 523–532.

